# Single molecule methylation profiles of cell-free DNA in cancer with nanopore sequencing

**DOI:** 10.1101/2022.06.22.497080

**Authors:** Billy T. Lau, Alison Almeda, Marie Schauer, Madeline McNamara, Xiangqi Bai, Qingxi Meng, Mira Partha, Susan M. Grimes, HoJoon Lee, Gregory M. Heestand, Hanlee P. Ji

**Affiliations:** Division of Oncology, Department of Medicine, Stanford School of Medicine, Stanford CA; Department of Electrical Engineering, Stanford University, Stanford CA

**Keywords:** methylation, single molecule sequencing, nanopore, cell-free DNA

## Abstract

Epigenetic characterization of cell-free DNA **(cfDNA)** is an emerging approach for characterizing diseases. We developed a strategy using nanopore-based single-molecule sequencing to measure cfDNA methylomes. This approach generated up to hundreds of millions of reads per cfDNA sample from cancer patients, an order of magnitude improvement over existing nanopore methods. Leveraging methylomes of matched tumors and immune cells, we characterized cfDNA methylomes of cancer patients for longitudinal monitoring during treatment.

## MAIN

Malignant tumor cells shed their DNA into the bloodstream of cancer patients in the form of cell-free DNA **(cfDNA)**. Sequencing cfDNA can identify cancer-associated biomarkers and is useful for disease monitoring. This approach is commonly referred to as a liquid biopsy^1-3^. Epigenetic modifications of tumor DNA are of particular interest because of their contribution to cancer development and progression^4^. Characterizing cancer-specific methylation changes has proven to be a highly sensitive and specific modality for liquid biopsies^5-7^. For detecting methylation, cfDNA is typically processed with bisulfite or enzymatic conversion of unmodified cytosines into uracils. Short-read sequencing detects the presence of methylated bases. However, this approach introduces biases such as significant GC skews, oxidative DNA damage, PCR amplification bias, and alignment artifacts^8,9^. Compounding these issues, extracted cfDNA from plasma has low yields. Characterizing cfDNA methylomes from patients remains challenging, particularly with conventional sequencing approaches.

Addressing these challenges, we developed a single-molecule sequencing approach for efficiently characterizing methylation profiles from the cfDNA of cancer patients **(Fig. 1A)**. This PCR-free process generates sequencing libraries from nanogram amounts or less of cfDNA per sample. We leveraged the Oxford Nanopore platform to identify cfDNA methylation without cytosine conversion. The passage of methylated DNA through the nanopore generates a unique electrical signal compared to unmodified DNA; currently, 5-methylcytosine **(5mC)** CpG methylation is detected with machine learning algorithms at high accuracy^10,11^. By eliminating PCR, we avoided GC-biased amplification skews from bisulfite conversion and enabled direct single-molecule counting of cfDNA. The nanopore-based methylation profiles thus directly reflect the native single-molecule state of the cfDNA.

**Figure 1.**
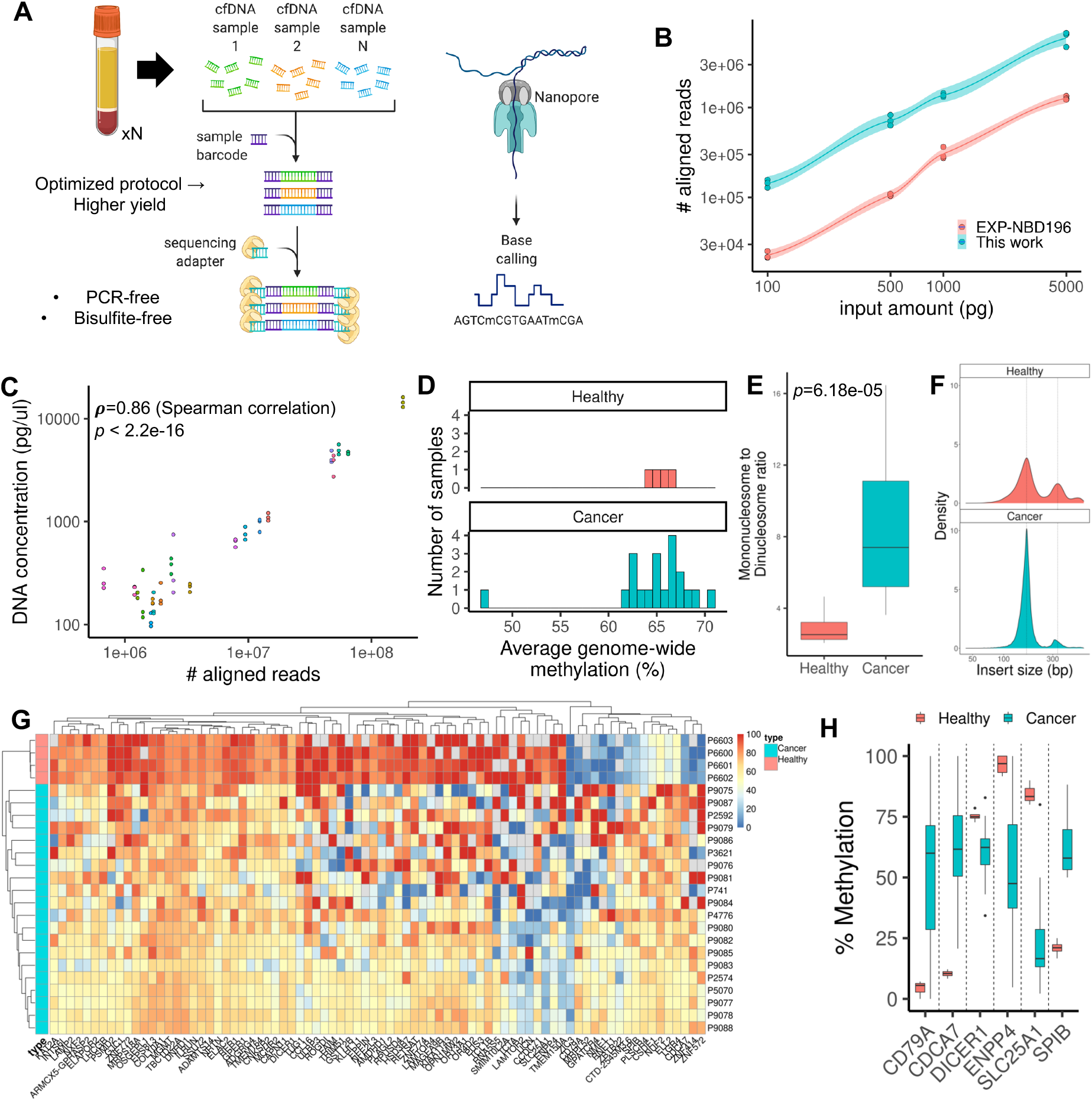
Nanopore sequencing of cfDNA. **(A)** An optimized protocol for generating cfDNA sequencing libraries enables high-throughput methylation characterization. **(B) Cell free DNA library comparison**. An optimized workflow enables approximately an order of magnitude increase in sequencing yield versus the conventional protocol. **(C) Sequencing yield correlation with input cfDNA**. Fluorometric quantification was performed on cancer patient-derived cfDNA samples and compared to the aligned sequencing yield. Correlation and significance value are annotated on the plot. **(D) Genome-wide methylation quantification**. The degree of methylation across the genome was computed for healthy and patient-derived cfDNA. **(E) Nucleosome enrichment analysis**. The ratio of mono-nucleosomes to di-nucleosomes was quantified for each tissue type, using a cutoff of 250bp between mono- and di-nucleosomes. **(F) Distribution of fragment sizes**. Example fragment sizes are shown for healthy and patient-derived cfDNA. Mono- and di-nucleosome size peaks are annotated with dotted lines to be 167bp and 334bp. **(G) Methylation profiles of healthy-and patient-derived cfDNA**. Gene-level methylation values for each sample were determined, and statistically significant ones (q<0.01) are plotted as a heatmap with the gene-level methylation percentage as the intensities. The heatmap was clustered by gene-level methylation. **(H) Differential methylation**. Statistically significant differences in methylation between sample types are shown for several selected genes.

Nanopore sequencing typically requires hundreds of nanograms of DNA for library preparation. However, extracted cfDNA yields range from single nanograms or less per ml of plasma. PCR amplification erases DNA methylation and cannot be used for nanopore-based methylation analysis. To enable PCR-free library preparation from nanogram DNA amounts of cfDNA, we identified a series of steps to efficiently incorporate sample barcodes and nanopore sequencing adapters to cfDNA **(Supplementary Figure 1)**. We systematically optimized reaction conditions to maximize cfDNA library yield. The method also incorporated a second end-repair and a-tailing step to add newer sequencing adapters for multiplexing cfDNA samples **(Methods)**.

To validate our approach, we developed a model DNA analyte that replicates some of the fragmentation patterns of cfDNA. We performed DNAse digestion on nuclei from isolated peripheral blood mononuclear cells **(PBMCs)**, where digestion of open chromatin yields DNA fragmentation patterns with a mononucleosome peak as seen in cfDNA **(Supplementary Figure 2)**. With this DNA, we used our library preparation method to generate libraries that were sequenced on the Oxford Nanopore PromethION system. We used the ‘megalodon’ software package, previously benchmarked to provide high-quality methylation calls from nanopore sequence data^10,11^, for sequence alignment and methylation calling **(Supplementary Figure 3)**. We sequenced libraries made from different amounts of PBMC nucleosomal DNA. At 5 ng of input DNA, the yield was approximately 6 million aligned reads inclusive of quality score filtering. With 100pg of input DNA, the yield was approximately 140,000 aligned reads **(Supplementary Table 1)**. We also prepared sequencing libraries of the same DNA with a standard protocol from Oxford Nanopore Technologies. Our method improved the aligned read yield by approximately an order of magnitude, even with input amounts as low as 100pg **(Fig. 1B)**.

Next, we sequenced cfDNA from 20 patients with colorectal cancer **(Supplementary Table 1, 2)**. Sequence yields ranged from one to 180 million reads per sample. We used a fluorometric assay to quantify the cfDNA of each sample **(Fig. 1C)**; the measurements were highly correlated with the total sequencing yield (Spearman’s rho = 0.86, *p* < 2.2e-16). As a control, we sequenced cfDNA from healthy individuals. There were several significant differences when comparing healthy and cancer patient cfDNA. First, the overall variance in genome-wide methylation in cancer patient cfDNA samples was higher at 7% compared to less than 2% in healthy cfDNA **(Fig. 1D)**. The variation may be indicative of aggregate methylation shifts due to an increase in tumor-specific cfDNA in plasma.

We distinguished mono- and di-nucleosome fragments using a size cutoff of 250bp **(Fig. 1E, F, Supplementary Figure 4)**. We observed that cancer patient cfDNA was enriched in mononucleosomes by approximately a factor of two (*p*=8.814e-05) compared to healthy controls. A similar result was also reported in another study^12^.

We determined the extent of gene-level methylation **(Methods)**. We observed many significant differences in gene-level methylation (*q* < 0.01) when comparing healthy versus patient cfDNA **(Fig. 1G, H, Supplementary Table 3)**. For example, there was an increase in the methylation of an immunologic marker gene *CD79A*^*13*^, a decrease in methylation of a tumorigenic modulation gene *DICER1*^*14*^, and a decrease in methylation of *SLC25A1*, a critical gene in mitochondrial homeostasis that is highly upregulated in cancer^15,16^.

We further examined the genes with statistically significant methylation differences between healthy controls and patient cfDNA. We calculated that such differences were not strongly correlated (Spearman’s rho = −0.188, *p*=7.392e-16) with the number of CpG sites covered per gene **(Supplementary Fig. 5)**. This result indicated that read yields were not a major confounding factor in our analysis. We also observed that the overall variation in gene-level methylation for these statistically significant genes was comparatively smaller in healthy cfDNA samples. This result indicated a relatively uniform cfDNA methylome across these samples **(*p*=4.57e-4, Supplementary Fig. 6)**. Finally, enrichment analysis of genes with statistically significant methylation differences using EnrichR^17,18^ yielded hits in the Myc pathway **(Supplementary Figure 7)**, suggesting that the changes in gene-level methylation were cancer-specific.

We determined whether individual reads from cfDNA sequence data can be classified as originating from tumor or immune cells. We leveraged patient-matched samples, which included resected tumors, peripheral leukocytes, and blood samples taken during treatment **(Fig. 2A)**. DNA was extracted from these matched samples and underwent nanopore sequencing with methylation profiling. To classify each read, we calculated the proportion of matching methylation sites based on genomic coordinate and methylation states when compared to the matched tumor or immune cell methylation profile. The result is a classification score for each read **(Supplementary Figure 8)**. After normalization, thresholds were used to classify reads as immune cell-or tumor-derived **(Methods, Supplementary Figure 9)**.

**Figure 2.**
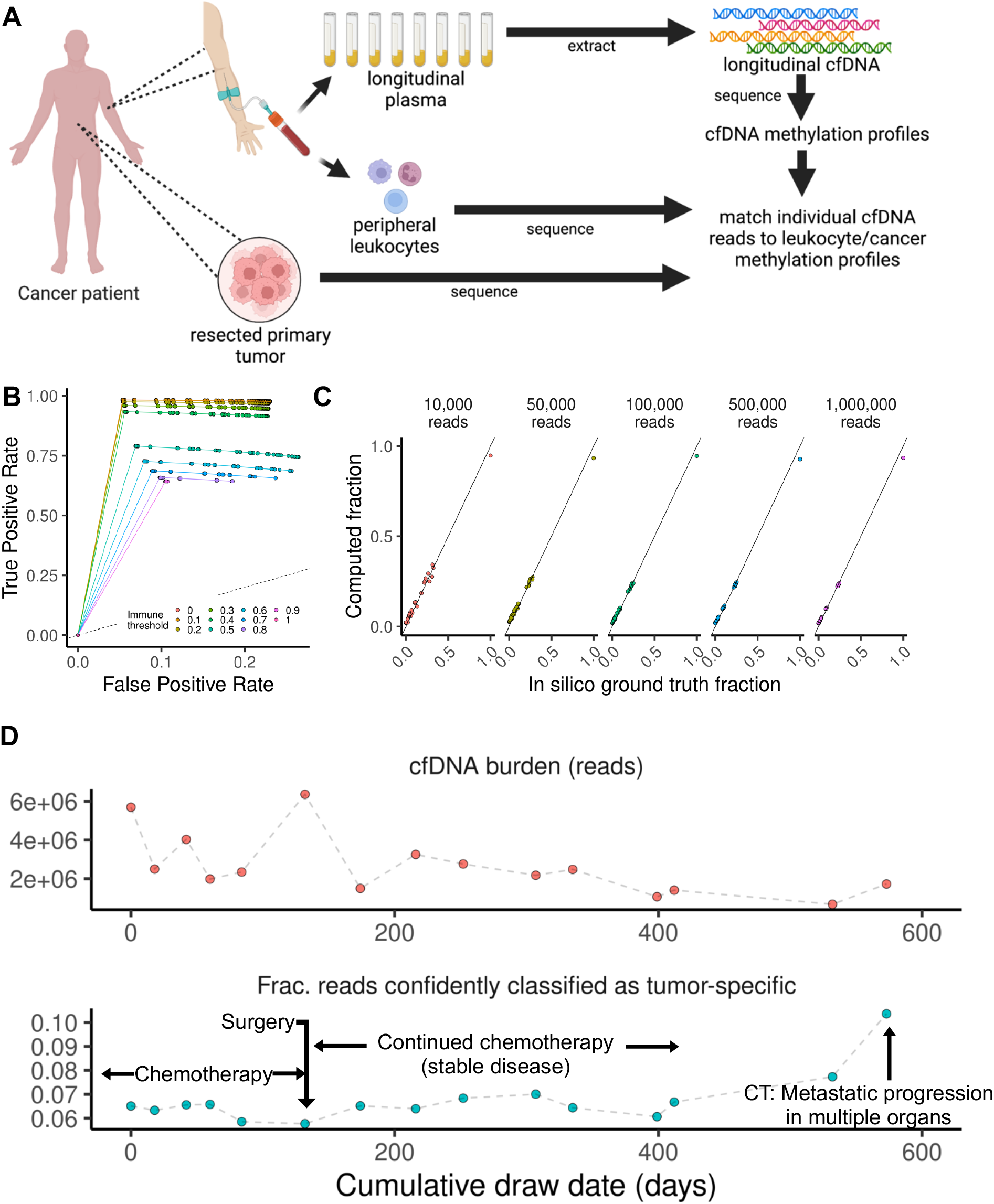
Single-molecule methylated sequence classification. **(A) Overview of method**. For a given patient, matched primary tumor tissue and peripheral leukocytes were obtained as reference samples alongside longitudinal plasma samples. Methylation data from the cfDNA is then classified leveraging the methylation profile of the reference samples. **(B) Classification accuracy**. We used GP2D and healthy donor-derived nucleosome mixtures to validate the classification procedure. ROC curves are plotted, where each curve represents a distinct immune threshold score. The curve is plotted by varying the cancer threshold score. **(C) Admixture validation**. The proportion of reads classified as belonging to cell line reference is plotted as a function of the actual admixture ratio and sequencing depth. **(D) Longitudinal methylation profiles of patient-derived cfDNA in colorectal cancer**. The overall cfDNA sequencing yield (upper panel) is plotted against the number of reads with methylation profiles matching that of the matched tumor with a calculated score of >0.9 (lower panel). Clinically relevant events are annotated. CT refers to computed tomography imaging.

We validated this approach using *in silico* admixtures between digested PBMC nucleosomes from a healthy donor and the GP2D cancer cell line. Nucleosomal DNA was subject to high-depth nanopore sequencing and methylation calling. Admixtures were computationally generated, and their proportions were estimated using our classification approach **(Methods)**. We measured the classification accuracy against the ground truth based on the source of any given read **(Fig. 2B)**. Performance was up to 97% sensitivity and specificity of 93% with an AUC of 0.951 when using stringent threshold cutoffs **(immune threshold: <0.1, cancer threshold: >0.9, Supplementary Figure 10)**. There was a corresponding trade-off with a declining proportion of classified reads **(Supplementary Figure 11)**; reads that could not be confidently classified as either type would be excluded from further analysis. We also simulated varying number of reads and did not observe declines in quantification performance when using stringent cutoffs **(Fig. 2C)**.

We also generated experimental admixtures where GP2D nucleosome DNA was added to donor nucleosome DNA while also varying the total quantity of DNA in the reaction. We observed a corresponding increase in cancer-derived reads at higher GP2D admixture fractions with Pearson correlation coefficients, ranging from 0.85 to 0.96 depending on the input amount **(Supplementary Figure 12)**.

For three patients (P4822, P6199, P6527) with different gastrointestinal cancers, we analyzed cfDNA from a longitudinal series of blood samples. The methylation profiles revealed responses to specific treatments and the emergence of treatment-resistant metastatic cancer. We performed nanopore sequencing on the set of patient-matched tumor, peripheral blood, and longitudinal plasma samples and determined their methylation profiles **(Fig. 2A, Supplementary Table 1**,**2)**. Matched tumor and immune cells were sequenced up to 28x coverage. Their methylomes, intersected by genomic position and filtered by coverage, yielded tens of millions of CpG sites per patient.

We observed longitudinal trends that correlated with specific clinical events based on our analysis. For one patient (P6199) receiving treatment for metastatic colorectal cancer, we sampled blood over approximately 600 days **(Fig. 2D)**. After having undergone chemotherapy and surgery, the patient had a period of stable disease. However, starting after day 400, the fraction of reads with tumor-specific methylation changes dramatically increased. This change correlated with CT imaging which showed substantial metastatic progression in multiple organs. Gene-level methylation analysis of this patient’s longitudinal cfDNA also showed gene-level methylation changes also found in the matched tumor, such as the Wnt/β-catenin regulator *TCIM*^*19*^ **(Supplementary Fig. 13)**.

We performed similar analyses on two other patients with metastatic cancer. One patient (P4822) had metastatic pancreatic neuroendocrine carcinoma and received multiple treatments, including targeted therapy, radiation, and peptide receptor radionuclide therapy **(PRRT)** spanning over 1,000 days **(Supplementary Fig. 14)**. After each treatment, there was a correlation between treatment effect and a drop in tumor-specific reads. The emergence of new metastases was reflected in a rise in reads with tumor-specific methylation changes. For another patient (P6527) with metastatic cholangiocarcinoma, resistance to the initial chemotherapy with gemcitabine was evident, but disease was reduced under a dual chemotherapy treatment **(Supplementary Fig. 15)**. The patient underwent extensive surgical resection of the primary tumor and liver metastases, as reflected in the immediate drop in tumor-specific reads. However, a subsequent rise in tumor-specific reads coincided with metastatic cancer recurrence.

We demonstrated a single-molecule approach for efficiently analyzing methylomes from cfDNA at scale. Improving on existing approaches by approximately an order of magnitude, our method yielded over 1 billion aligned reads across our cancer cohort when using small amounts of cfDNA. We detected cancer-associated methylation profiles with distinguishing epigenetic characteristics and provided a measurement of tumor burden that corresponded to clinical events. Our work demonstrated streamlined methylation analysis of cfDNA with significantly fewer experimental procedures and bottlenecks than short-read sequencing **(Supplementary Table 4)**. Newer machine learning models that incorporate the detection of other modified bases such as 5-hydroxymethylcytosine **(5hmC)** methylation can be applied to archived raw data. In summary, we describe a single-molecule sequencing method that enables the analysis of cfDNA methylation. This approach has the potential to impact liquid biopsy diagnostics for cancer detection and characterization.

## METHODS

### Samples

We obtained informed consent from all patients based on a protocol approved by Stanford University’s Institutional Review Board. Blood and tissue samples came from the Stanford Cancer Center, the Stanford Tissue Bank, and the Stanford Blood Center. For a subset of patients enrolled at the Stanford Cancer Center, we obtained whole blood samples in Streck or EDTA tubes which were later centrifuged into plasma and PBMC fractions. For these patients, matched tumor tissue was also obtained. Plasma from the Stanford Tissue Bank was obtained as single aliquots in 1ml cryovials. Tumor tissue was archived by flash freezing in liquid nitrogen and stored at −80C. From the Stanford Blood Center, we obtained whole blood from anonymous donors to serve as healthy controls; these samples were centrifuged into plasma and buffy coat fractions. All samples were stored at −80C before processing.

### Nucleosome DNA controls

To generate DNA fragments modeling the qualities of cell-free DNA, we used the EZ Nucleosomal DNA Prep Kit (Zymo Research). This method uses DNAse to digest open chromatin positions and yields a fragment pattern characteristic of cell-free DNA instead of random fragmentation. Briefly, nuclei were processed from whole cells by adding a nuclei prep buffer that lyses the cell membrane but leaves the nuclei membrane intact. Enzymatic DNAse digestion then fragments DNA at unprotected locations, after which DNA is purified with the kit’s included components. For nucleosomes from cancer lines, we used cells treated with trypsin.

The Stanford Blood Center provided anonymous donor blood as healthy controls. We used peripheral blood mononuclear cells (PBMCs) for nucleosome preparation. Whole blood was diluted with an equal volume of PBS and added to a SepMate PBMC isolation tube (STEMCELL Technologies) containing Ficoll. The tube was spun at 1200g for 10 minutes before decanting into a new tube. Cells were spun again at 400g for 5 minutes and were washed with PBS. Cells were resuspended in freezing medium (90% FBS/10% DMSO). Isolated PBMCs were then used as input for the nucleosome preparation kit. For the experimental admixtures, the PBMC and cell line nucleosomes were diluted to a target concentration (e.g. 1ng/ul) and mixed to known ratios. Serial dilutions of this mixture are then performed to simulate lower input amounts.

### Sample processing

Extracted DNA was obtained from tissue biopsies using the Maxwell 16 DNA extraction kit (Promega). Briefly, a small tissue fragment was excised from the tissue sample with a scalpel and deposited into the input well of the DNA purification cartridge. The cartridge was placed into the Maxwell 16 instrument (Promega), and the associated protocol was run. For extracting cell-free DNA, plasma was separated from whole blood by centrifugation. The plasma fraction was pipetted into a Maxwell 16 ccfDNA Plasma kit cartridge (Promega) using the standard instrument protocol. The cellular blood portion was extracted using a Maxwell 16 LEV Blood DNA Kit. Yields were measured by Qubit (Thermo Fisher Scientific). Cell-free DNA was quantified using the AccuBlue NextGen DNA Quantification Kit (Biotium).

### Sequencing library preparation

We developed a protocol for generating sequencing libraries that accommodate the low input amounts of cfDNA and maximize sample barcode adapters’ incorporation rate. Briefly, 25ul of extracted cfDNA (out of a typical 50ul extracted volume; thus corresponding to ∼0.5ml of plasma) was diluted with 25ul of water. The sample DNA underwent end-repair and A-tailing with conditions of 20C for 30 minutes and 65C for 30 minutes (Roche KAPA HyperPrep kit). We ligated native barcodes using 5ul of each barcoded adapter (EXP-NBD196, Oxford Nanopore Technologies) following the standard reaction volumes in the KAPA HyperPrep workflow. We used a thermocycler for the ligation step for 4.5 hour incubation at 20C before holding at 4C overnight to maximize the ligation yield. These steps provided a higher ligation rate of cell-free DNA molecules to a native barcode adapter than the standard protocol’s shorter end-repair/A-tailing and ligation time (10 minutes per the standard Oxford Nanopore protocol).

After the ligation step, 88ul of Mag-Bind Total NGS beads (Omega Bio-Tek; an alternative to Ampure XP beads) were added and mixed to each reaction. After incubation for 5 minutes, the mixtures were pooled together into a 50ul centrifuge tube. The beads were magnetized and washed with 80% ethanol using a DynaMag separation rack (Thermo Fisher Scientific) before eluting in 600ul of 10mM Tris-HCl pH 8.0 buffer. We performed a second bead cleanup step with 900ul Mag-Bind Total NGS beads (1.5X ratio) and the same magnetic rack procedure. The elution solution was 50ul 10mM Tris-HCl pH 8.0 buffer.

We improved the preparation of multiplex sequencing libraries. For the Oxford Nanopore platform, multiplexing is restricted to the AMII adapter, which has the same motor protein family as the LSK109 sequencing chemistry. This adapter has a significant disadvantage for short fragment libraries because it incurs active consumption of on-chip “fuel” of idle sequencing molecules, leading to rapid flow cell exhaustion. To address this issue, we modified the library preparation process to incorporate the updated “fuel-fix” adapter (LSK110 kit, Oxford Nanopore Technologies), which at the time when these experiments were conducted, did not have multiplexing capabilities. We developed a protocol to enable multiplexing with this adapter. We performed a second end-repair and A-tailing reaction using the Kapa HyperPrep library preparation kit. This step removed the sticky end from the barcode multiplexing adapter and produced a compatible A-tail for sequencing adapter ligation. We used an increased amount (10ul) of the AMX-F sequencing adapter (LSK110, Oxford Nanopore Technologies) for the ligation step to maximize the yield of sequencing adapters to barcoded fragments. This second ligation reaction occurred for 1.5 hours. Subsequently, we mixed in 88ul of Mag-Bind Total NGS beads and incubated for 5 minutes. As in the standard protocol, we washed the beads with 200ul SFB buffer (Oxford Nanopore Technologies) with gentle tube flicking to resuspend the beads during the wash steps. The beads were resuspended in EB buffer (Oxford Nanopore Technologies). We used 1ul for quantification with Qubit (Oxford Nanopore Technologies) and 1ul for determining the DNA size with an E-gel EX cartridge (Thermo Fisher Scientific).

We generated sequencing libraries for tumor tissue and PBMCs with 1-2ug of extracted genomic DNA. For some tissue and buffy coat samples with low extraction yields (less than 1ug), we used the entire amount of extracted DNA for library preparation. We followed the standard Kapa HyperPrep library preparation kit protocol using 5ul of AMX-F adapter (LSK110) without barcoding. Each sample was loaded into its own PromethION flow cell for sequencing. For comparison with the standard library preparation protocol, we followed the standard protocol for Native Barcoding (EXP-NBD196) coupled with the SQK-LSK109 library preparation kit using the AMII adapter. The standard protocol is available on the Oxford Nanopore Technologies website.

### Nanopore sequencing and data processing

We performed sequencing on the Oxford Nanopore Technologies’ PromethION 24 instrument. Approximately 150fmol of the library was loaded for each flow cell, loading up to four flow cells in a single batch. The remainder of the library was stored at −80C. For tissue samples, we used one entire flow cell per sample. Sequencing runs had a duration of 72 hours. Barcode demultiplexing was performed on the sequencer using onboard basecalling in MinKNOW with the “high accuracy” model and then transferred to a separate storage device. Raw demultiplexed fast5 sequencing data were processed using Megalodon v2.4.0 (Oxford Nanopore Technologies, https://github.com/nanoporetech/megalodon) and Guppy v5.0.16 (Oxford Nanopore Technologies, available closed source at https://nanoporetech.com/) with the “dna_r9.4.1_450bps_modbases_5mc_hac_prom.cfg” model for each demultiplexed barcode folder with standard settings. The quality score cutoff was 7. The GRCh38 reference was used for alignment. The output consists of a file in BedMethyl format for each sample. The files included modified base calls, a sequencing alignment bam file with modified base calls for each read, and a per-read text file containing modified base call probabilities. Before further processing, the BedMethyl and sequence-alignment bam files were sorted and indexed with samtools^20^. For larger sequencing runs involving multiple samples (e.g., from multiple flow cells and many barcodes), data was transferred to the Sherlock High-Performance Computing cluster at Stanford University for multi-node GPU-based data processing.

The overall methylation status of sequenced cfDNA was determined by taking the average of all methylation values across all sequenced sites that had at least one read (coverage > 0). To determine nucleosome enrichment, we subsampled each library’s sequence aligned bam file to 50,000 reads, tabulated the estimated fragment size as inferred by the alignment length, and set a cutoff of 250.5 base pairs separating mono- and di-nucleosome states, with a maximum length filter of 600bp. This data was then compiled for all reads and all samples sequenced.

We determined gene-level methylation for all sequenced cfDNA samples by calculating the average per-site methylation for each CpG site with non-zero coverage, and then searched for statistically significant differences in gene-level methylation. This procedure is similar to that of another study^7^, with the main difference being that our data enables site-level detection of methylation percentage. We utilized “gene”-level annotations in GENCODE v38, which includes all coding exons and introns. First, we filtered on genes which were covered by at least one sample and where the standard deviation of average gene methylation is greater than zero. Then, grouping by gene-level annotations, we calculated the average methylation. Based on genomic coordinates, we then excluded annotations that were pseudogenes, unprocessed, “to be experimentally confirmed genes,” lncRNAs, and miRNAs. Finally, we used a t-test to compare methylation between the healthy donor-derived cfDNA and cancer patient-derived cfDNA. An FDR-based multiple testing correction was applied to determine statistically significant differences in gene-level methylation. We used a cutoff of q < 0.01.

### In silico admixture analysis

To simulate circulating tumor DNA **(ctDNA)** data of varying fractions in cfDNA, we generated *in silico* admixtures of sequence data from the GP2D cancer cell line-derived and PBMC-derived nucleosomes. Using a Python script, we mixed two sequence-aligned bam files using a known random seed to ensure reproducibility. We also controlled for the number of reads to simulate different read depths. Methylation profiles were compiled from the Mm and Ml tags using the modbampy library as part of the modbam2bed package (https://github.com/epi2me-labs/modbam2bed). We used only reads that mapped to the reference and used the subsequent bam file for downstream analysis. The remainders of the reads were not used, including unmapped reads and those with secondary or supplementary alignments. As another output, we included the metadata about the sample origins, namely whether it originated from PBMC-derived nucleosomes or a cancer cell line.

### Reference methylation profile processing of tumors and immune cells

For a subset of the patient samples, we had matched tumor tissue and PBMCs. These matched samples underwent nanopore sequencing to generate reference methylomes; methylation calls were also performed with megalodon. The reference methylomes consist of megalodon’s output BedMethyl files, which contain genomic positions of CpG sites, coverage (> 0), and the associated percent methylation for that position.

To process these reference profiles for read-level classification, we used an R script to read both the tumor and PBMC methylation profiles. We intersected these profiles on genomic coordinate positions, with a coverage filter of greater than four in both samples^21^. We considered a site to be methylated if the percentage methylation per a given genomic segment was greater than zero. The resultant intersected table was used for read-level classification (below).

To determine gene-level methylation for primary tumor and blood samples, average methylation profiles were determined for each “gene”-level annotation in GENCODE v38. These were then filtered to exclude annotations that were pseudogenes, unprocessed, “to be experimentally confirmed genes,” lncRNAs, miRNAs, miscRNAs, snRNAs, snoRNAs, scaRNAs, sRNAs, and rRNAs. We calculated the difference in methylation between the primary tumor and immune cells, and selected the top and bottom 25 genes. Using this gene list, we extracted the methylation values from the patient cohort that underwent longitudinal sampling.

### Single-molecule read classification to reference profiles

We built a computational workflow to classify whether an individual read is associated with an associated reference methylation profile. It consists of two steps

#### 1. Read-level methylation processing

This process utilizes sequence-aligned bam files containing read modifications (from megalodon). We used a python script to emit a table with columns consisting of the read name, genomic coordinate, and called methylation status. Thus this is a flat data table whereby genomic coordinates and their methylation states can be grouped by individual reads.

#### 2a. Scoring against reference profiles

We classified each read alongside a reference methylome containing informative methylation sites. Reference methylomes consist of matched tumor and immune cell methylation profiles that were nanopore sequenced and processed as above. Informative sites are CpG sites where the methylation values differed between sample types. This process generated a value 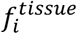 where *i* is the read number from 1 to the total number of aligned reads.

Specifically, 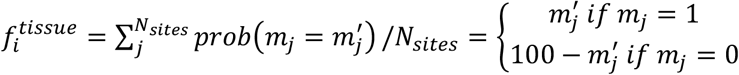 where *i* is the read number, *tissue* is the candidate reference profile to match against, *m*_*j*_ is the methylation of read *i* at site *j* (either 0 or 1), *m*_*j*_*’* is the methylation of *tissue* at site *j* (ranging from 0 to 100), and *N*_*sites*_ is the number of methylated sites to consider for read *i*.

In other words, 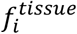 is the mean probability that read *i* matches to a specific *tissue* methylation reference profile. To implement this scheme, we obtained the methylation status (m_i_ … m_n_) of each CpG site for each read from a given sample and its reference coordinate. Then we intersected these coordinates of the CpG sites to the corresponding locations of a candidate tissue reference methylation profile (e.g., from PBMCs or matched primary tumor, with methylation profile m_i_’ … m_n_’). Subsequently, we calculated a matching score, where each site is scored m_i_’ if the m_i_ is methylated; otherwise, it is scored 100 - m_i_’. In other words, the score is the probability that the methylation site and value m_i_ is the same as the reference profile site m_i_’, which is equivalent to the reference profile’s methylation level at that site. It is then divided by the total number of candidate CpG sites on the read to derive 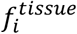. Reads with no candidate CpG sites or matching locations in a reference methylome were not considered.

#### 2b. Score normalization and thresholding

A normalized per-read tumor score was then assigned by the ratio of scores 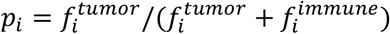,-with scores close to zero indicating likely matches to PBMCs, and scores close to one indicating likely matches to tumor tissue. A final classification is determined by setting thresholds for matching to PBMC and cancer methylation profiles. By using a dual threshold system, a subset of reads in between the thresholds do not have a confident/stringent classification and were not called to be either type (and thus can be neglected from the final analysis). We varied the two thresholds to determine ROC curves and AUC performance metrics.

#### 3. Processing all reads

We process and aggregate classification calls from all reads in order to calculate the fraction of reads with methylation changes that were classified as specific to the tumor.

## DATA AVAILABILITY

Aggregated methylation calls and sequence aligned BAM files are deposited in NCBI’s dbGaP under accession phs002950. Code is available at https://github.com/billytcl/nanopore_cfDNA.

## ACKNOWLEDGEMENTS

The authors acknowledge financial support from the Clayville Foundation (to H.P.J), National Cancer Institute (R33CA247700 to H.P.J.), and the National Human Genome Research Institute (R35HG011292 to B.T.L.). The authors also acknowledge Professor Brooke Howitt of the Stanford Tissue Bank, a Shared Resource Facility at the Stanford Cancer Institute, for providing banked frozen plasma for this study. The authors also acknowledge the Stanford Blood Center for providing fresh plasma from anonymous healthy donors for this study. Figures 1A and 2A were illustrated using BioRender.

## AUTHOR CONTRIBUTIONS

B.T.L, G.M.H., and H.P.J conceived the study. G.M.H. and H.P.J. established and oversaw the translational patient recruitment component of the study. A.A., M.S., M.M., processed the samples. B.T.L. performed the sequencing experiments. B.T.L., X.B., Q.M., M.P., S.M.G., H.L, and H.P.J. analyzed the data. B.T.L, and H.P.J. wrote the manuscript.

