## Supplementary Information for "Single molecule methylation profiles of cell-free DNA in cancer with nanopore sequencing"

### TITLE

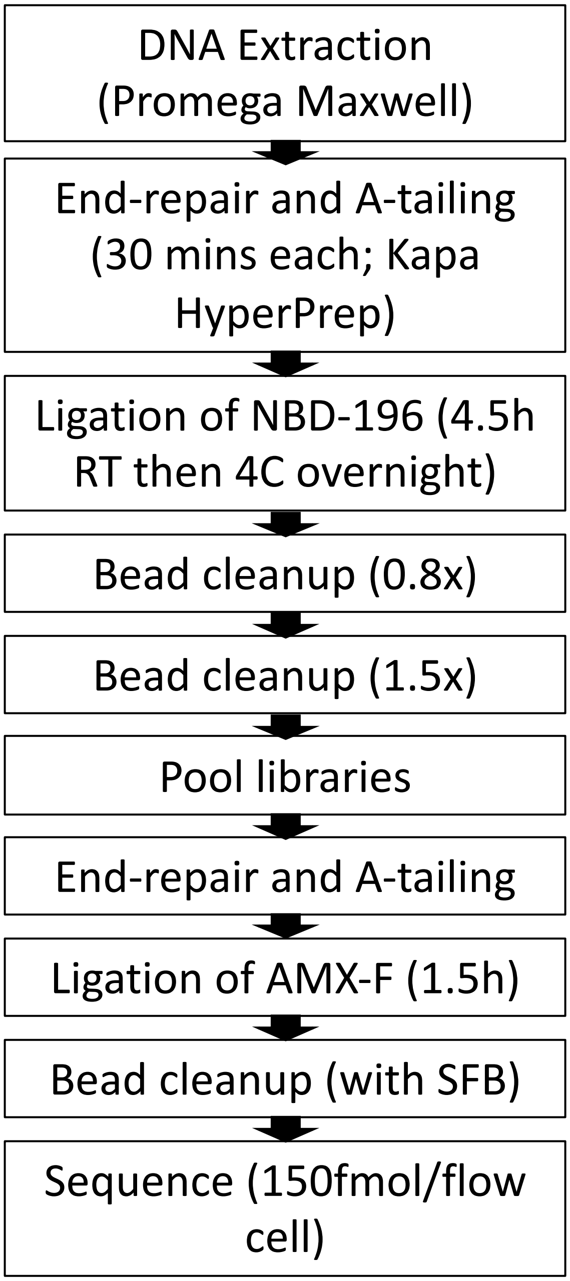


**Supplementary Figure 1. Sequencing library preparation workflow.** The library preparation workflow used in this study for cfDNA samples. Sequencing was conducted on the Oxford Nanopore platform. These steps maximized ligation yields versus standard protocols.

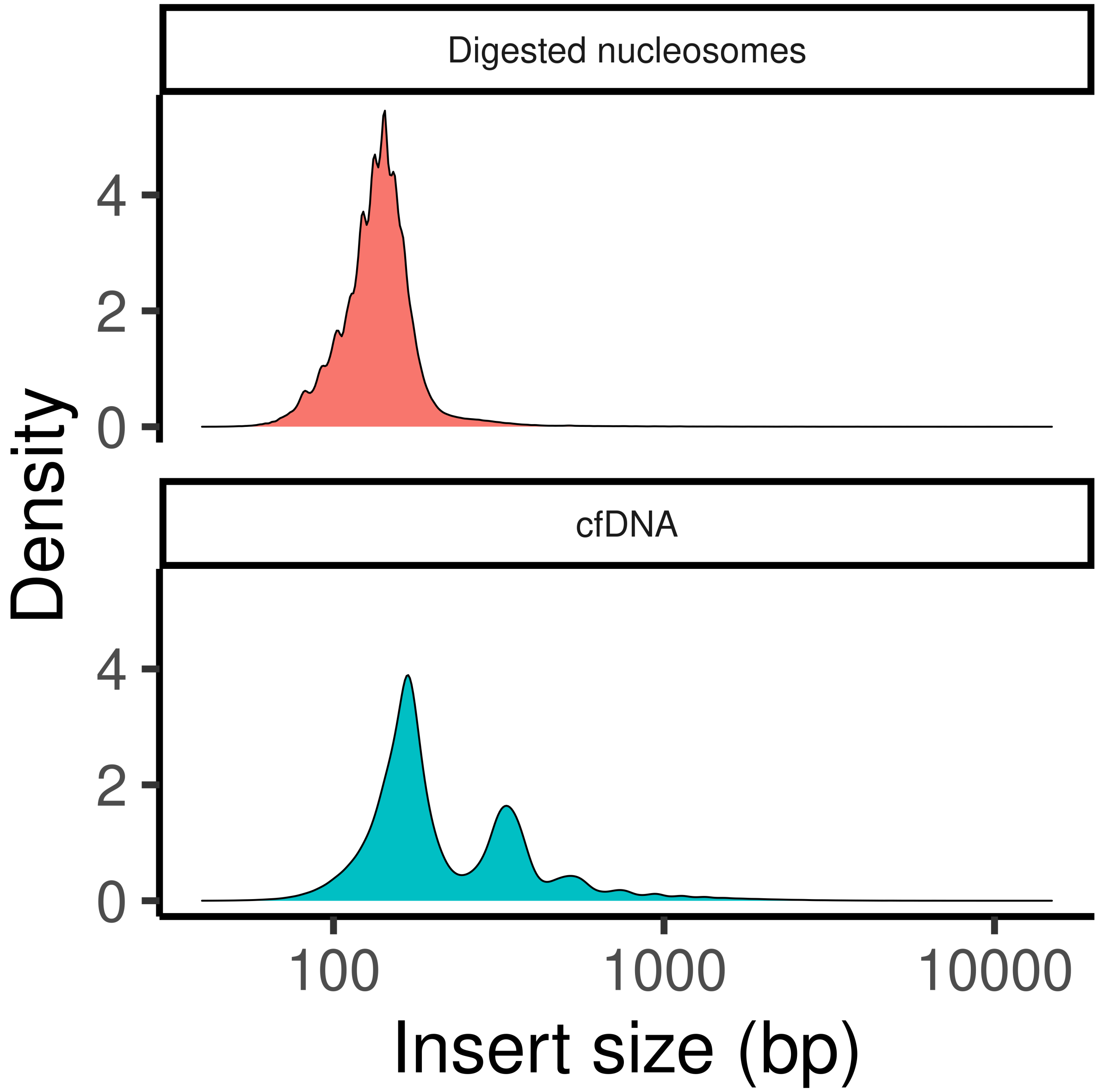


**Supplementary Figure 2. Digested nucleosome size comparison with cfDNA.** The insert size distribution of digested PBMC nucleosomes (top), which was used as a model analyte. This size distribution was compared to the size distribution of cfDNA. Secondary peaks in the cfDNA distribution correspond to dinucleosomes and higher sizes. The PBMC nucleosomes consisted only of mononucleosomes due to complete digestion of the open chromatin.

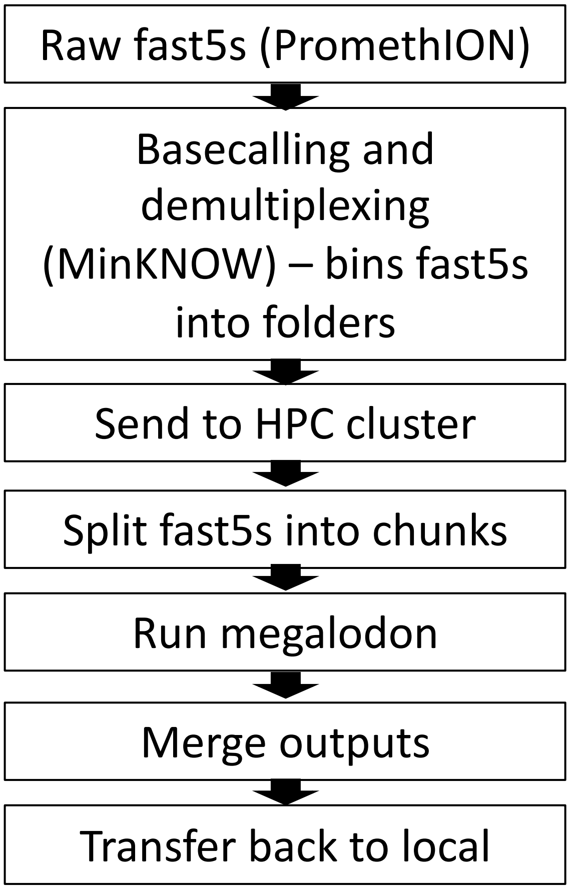


**Supplementary Figure 3. Computational workflow.** The workflow for calling methylation from nanopore-based cfDNA sequencing data is shown. These steps enable streamlined processing of large data volumes (>10TB).

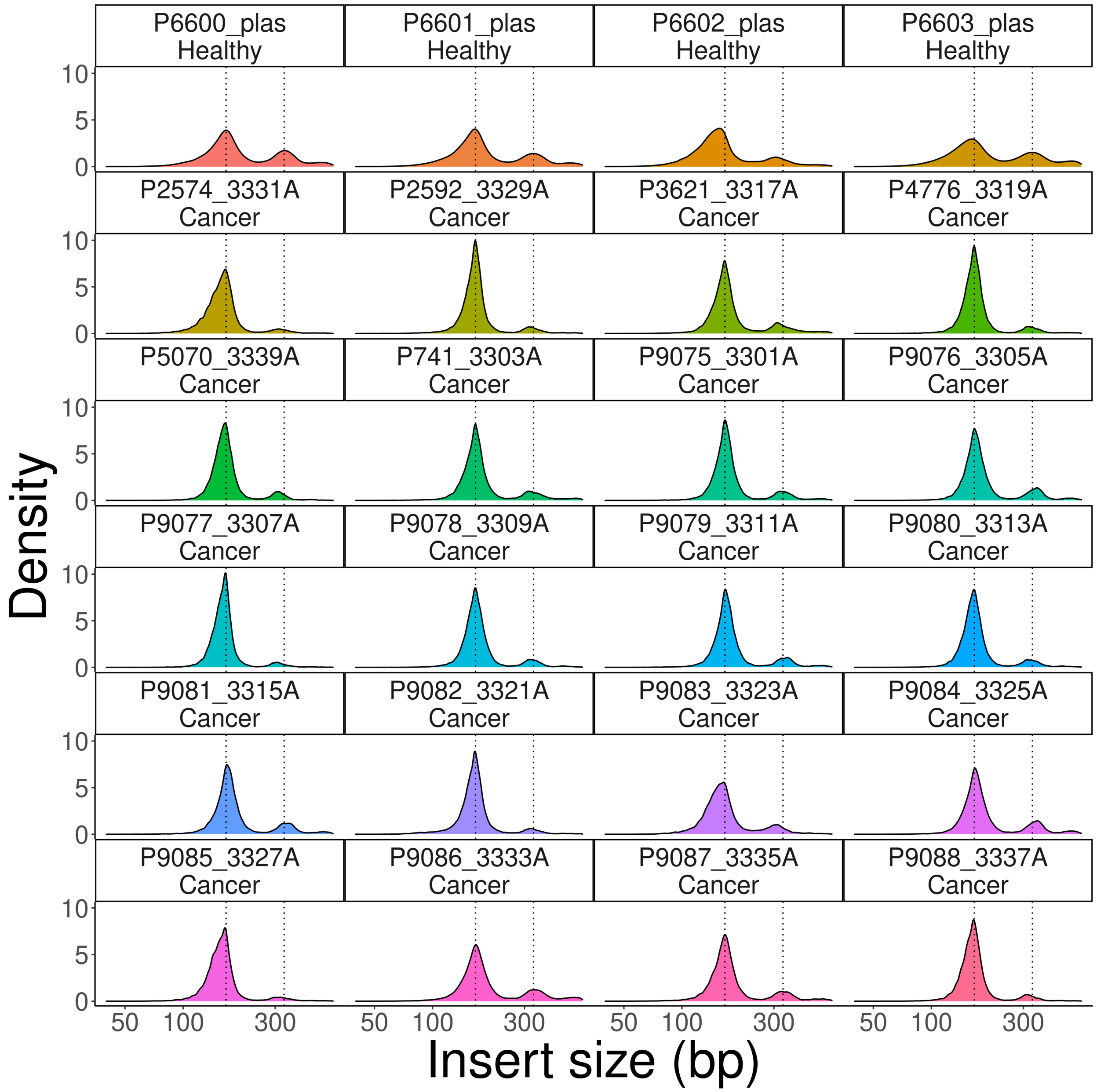


**Supplementary Figure 4. Fragment size distribution of healthy donor and cancer patient cfDNA.** The fragment size distribution of healthy control plasma and cancer patient-derived plasma is shown. The top row consists of cfDNA samples from healthy controls. The remaining rows come from cancer patients. Dotted lines indicated mono- and di-nucleosomes.

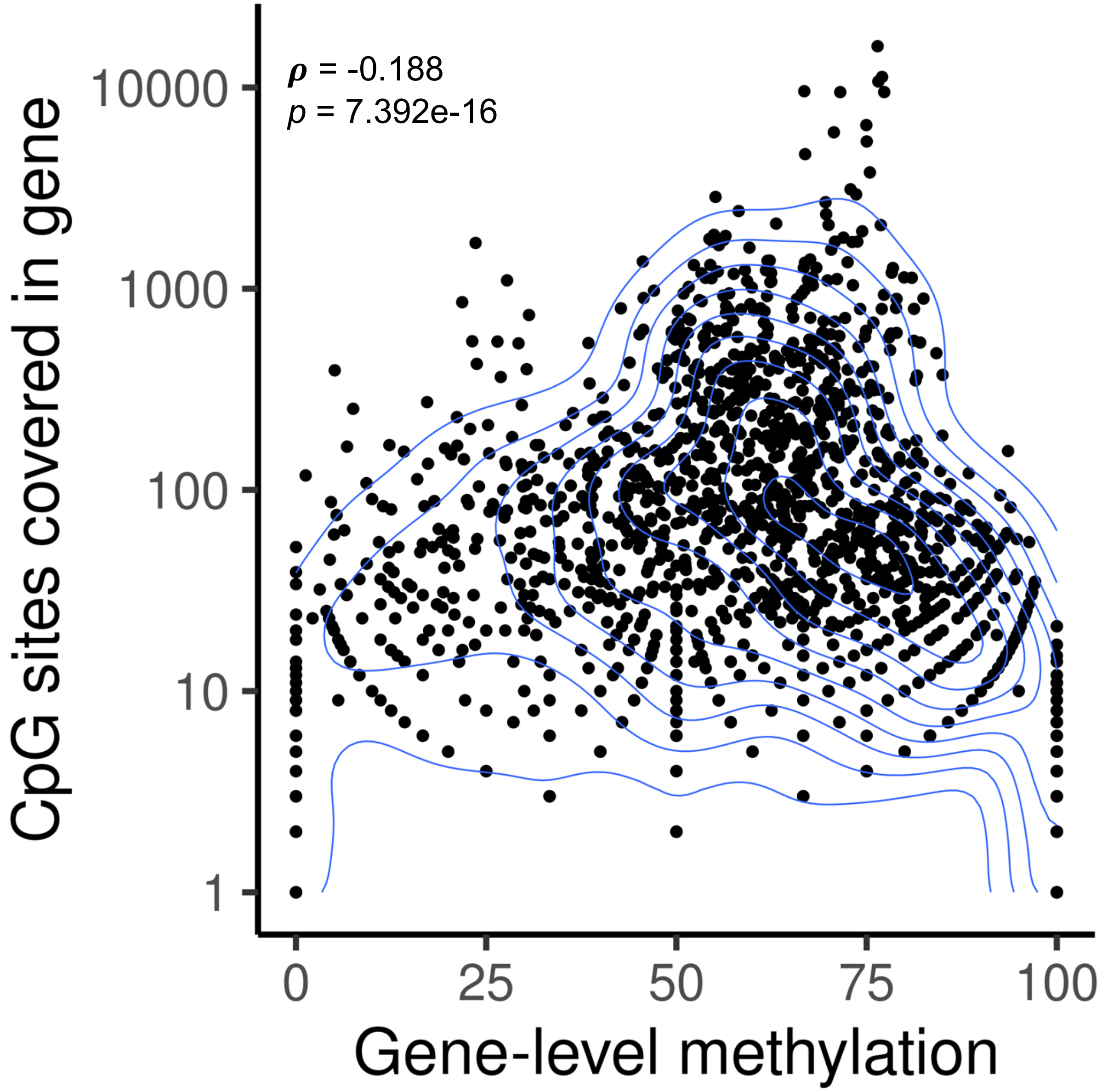


**Supplementary Figure 5. Correlation between CpG site coverage and gene-level methylation.** The number of CpG sites covered per gene and per patient is plotted against the calculated average methylation for that gene. A two-dimensional density contour plot is overlaid to show the density of scatter points. The Spearman correlation coefficient and associated p-value was also calculated and shown on the plot.

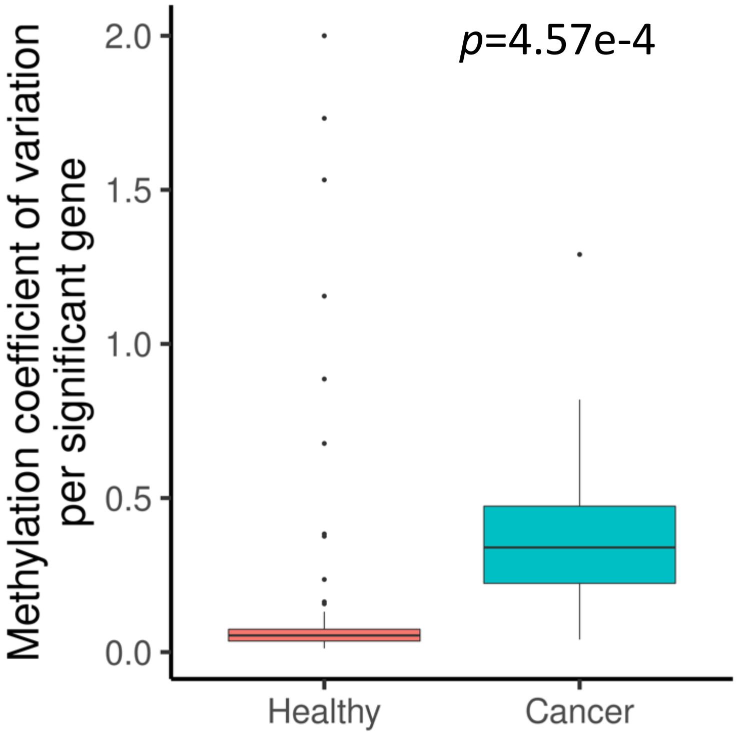


**Supplementary Figure 6. Variation in gene-level methylation between healthy and cancer profiles.** We identified genes with different levels of methylation when comparing cfDNA from cancer patient versus healthy controls. Genes that passed an FDR-based multiple-testing significance value of $q < 0.01$ were considered to have differential methylation. We calculated the coefficient of variation for the methylation values of each gene based on cancer patients versus controls.

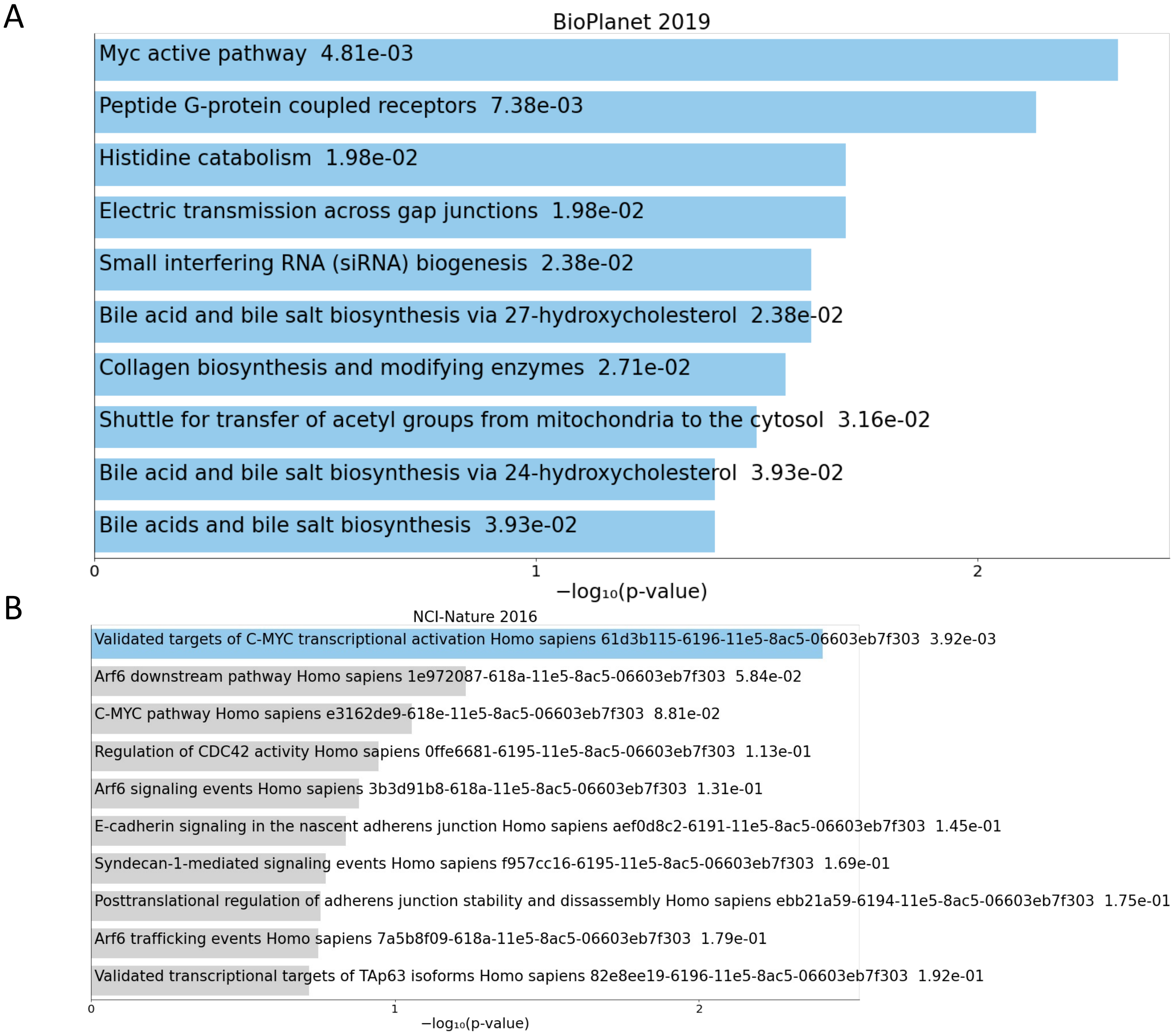


**Supplementary Figure 7. Gene enrichment analysis for cancer patient cohort.** Gene enrichment analysis was performed for significantly different genes between healthy and cancer patient-derived cfDNA. p-values are shown, alongside the associated pathway. Blue bars indicate $p<0.05$. A and B refer to two separate gene pathways curated by EnrichR.

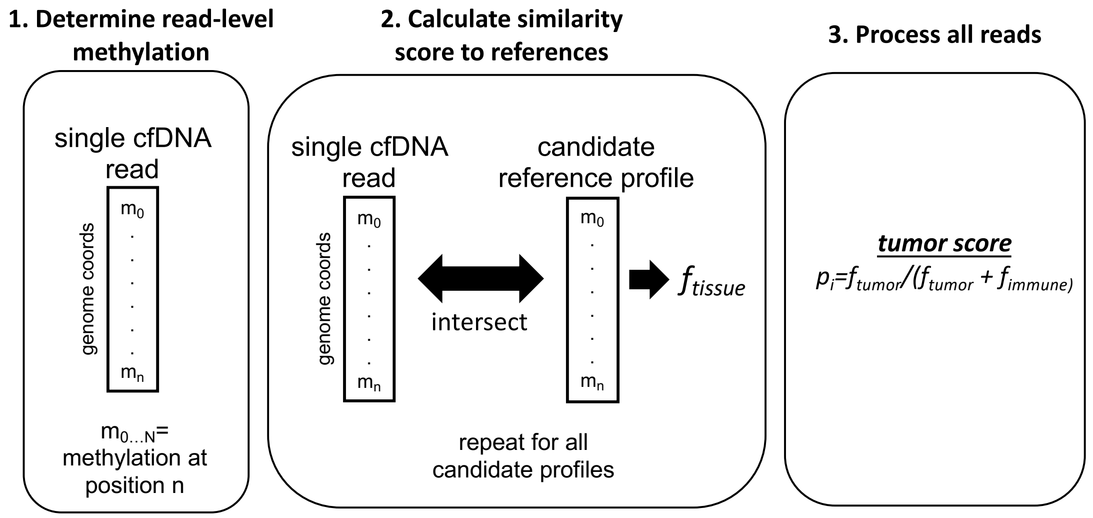


**Supplementary Figure 8. Framework for read classification.** Individual nanopore reads from cfDNA are classified by using reference profiles that come from matched tumor or PBMC/immune cell methylation data. Each read, their associated CpG sites, and their methylation states, are compared to candidate references. The calculated score reflects the similarity of a read to a particular candidate reference methylome. Regardless of their methylation status, all reads were processed with this framework.

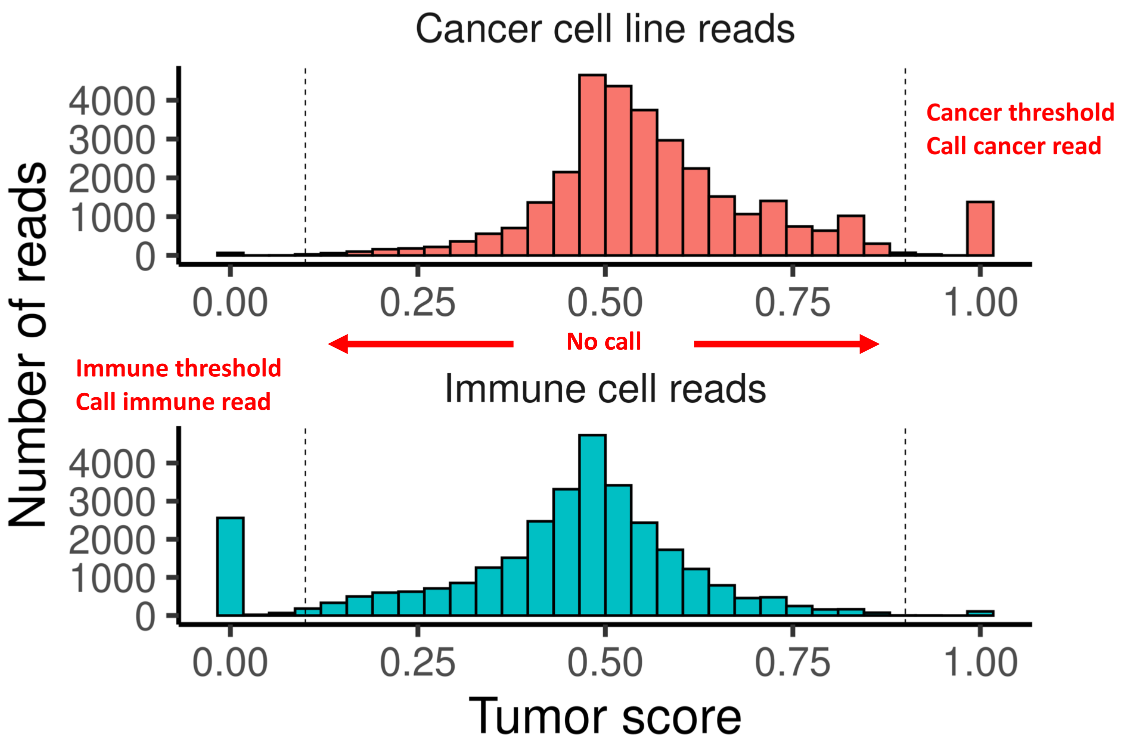


**Supplementary Figure 9. Distribution of tumor classification scores for single reads.** We provide an example of a tumor score histogram using an *in silico* admixture read data set. Each read has a calculated tumor score based on its methylation similarity to a matched tumor or immune reference profile. The title of each panel reflects the ground truth origin of each read set. Cancer reads are sequences that are mixed from GP2D cancer cell line nucleosomes that were nanopore sequenced and for which methylation calls were made, and immune cell reads are reads that are from healthy donor nucleosomes. There are two classification thresholds: one for immune cell origin, and one for classification of cancer cell origin. Reads matching the threshold criteria, such as tumor score > 0.9 or < 0.1, are classified as tumor-specific or immune-specific respectively. Reads falling outside the thresholds are not classified and are excluded from analysis.

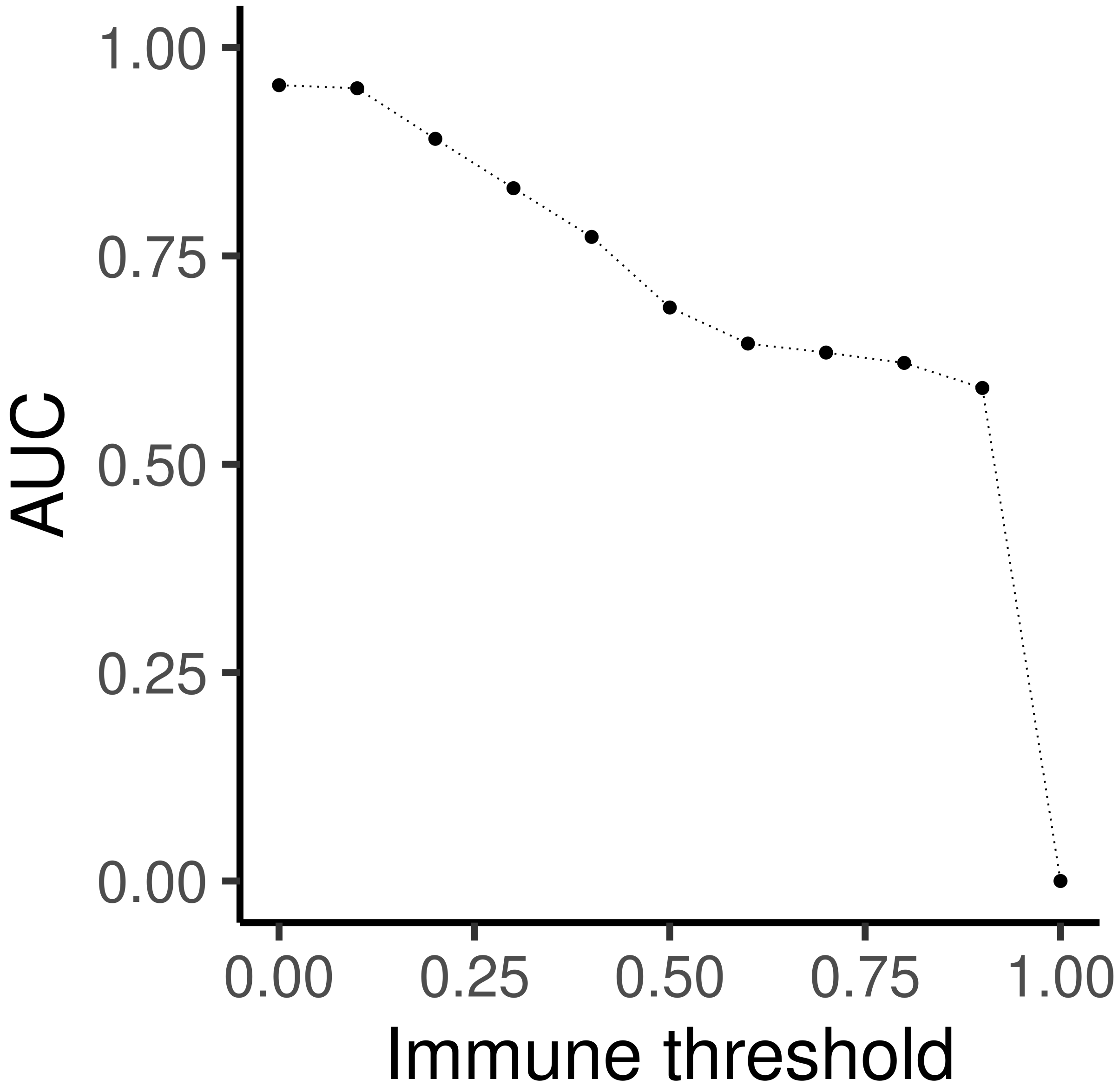


**Supplementary Figure 10. Classification AUC for various thresholds.** The AUC is calculated for various immune threshold values for one set of an *in silico* admixture between cancer cell line and healthy donor methylome data.

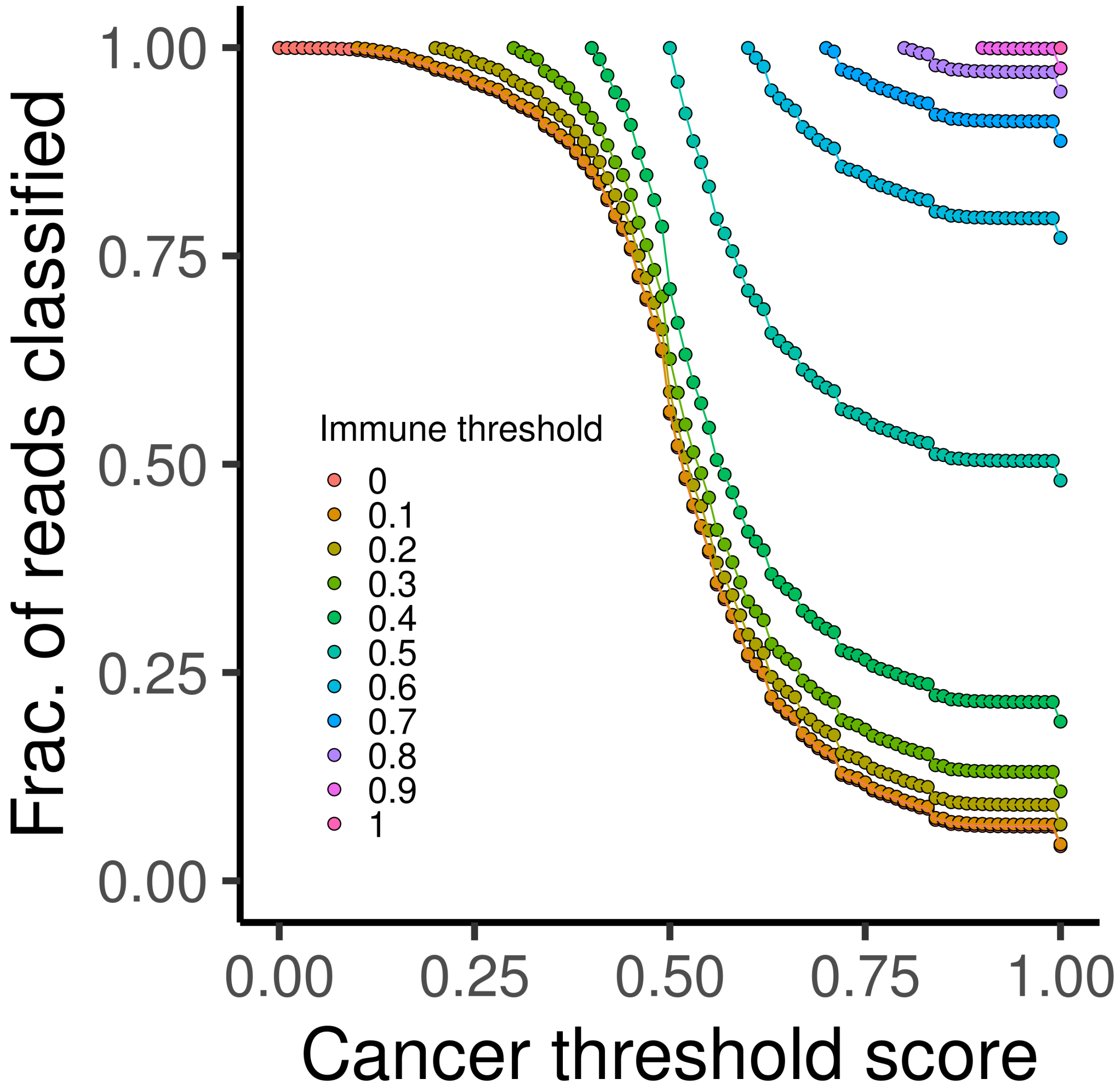


**Supplementary Figure 11. Sequence yield for read classification.** The sequence yield for reads being classified is shown for various immune threshold values for one set of an *in silico* mixture between cancer cell line and healthy donor methylome data.

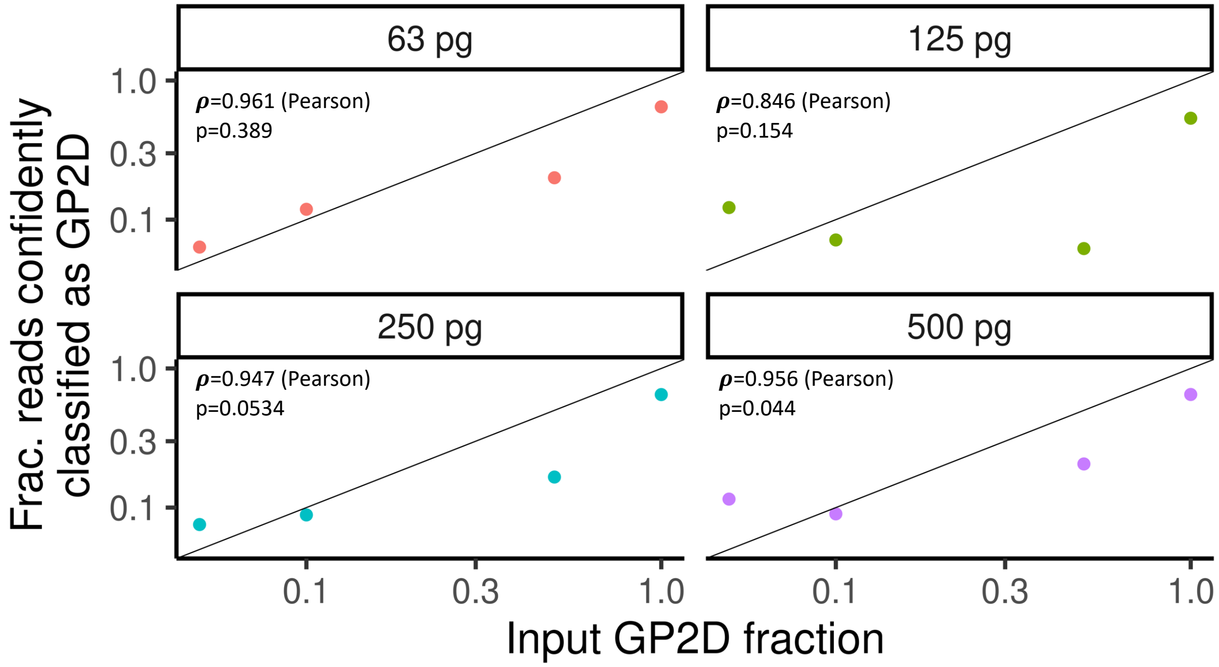


**Supplementary Figure 12. Experimental admixtures.** Experimental admixtures were performed between digested nucleosomes of the cancer cell line GP2D and healthy donor PBMCs. Various mixture fractions and input amounts are shown.

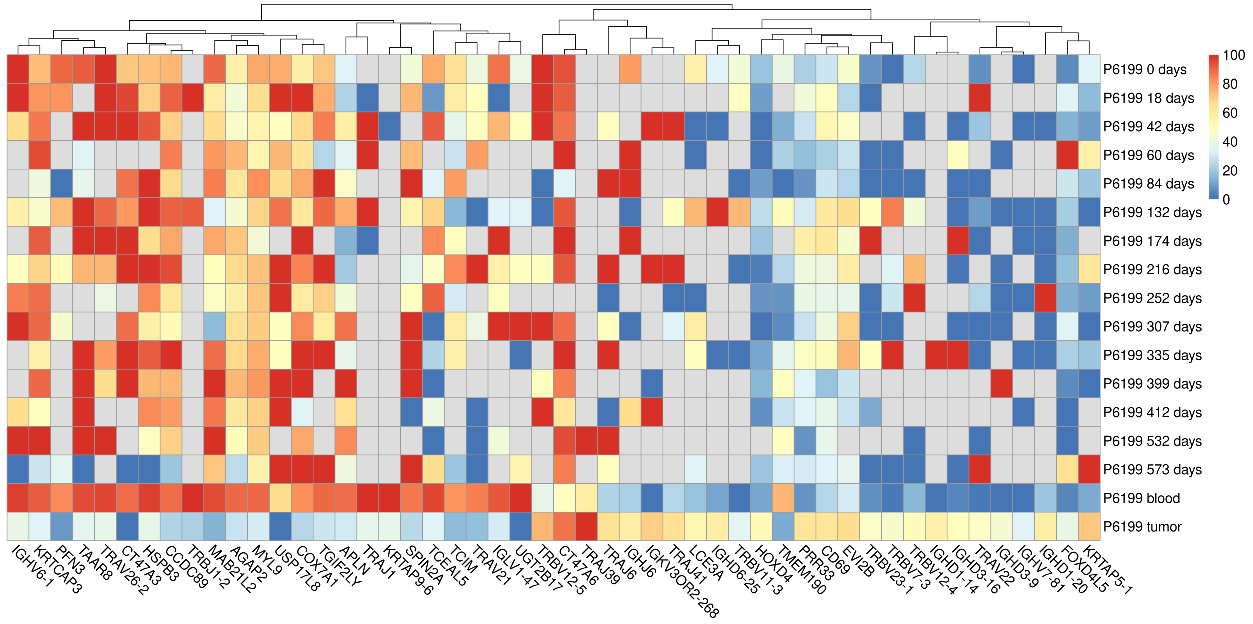


**Supplementary Figure 13. Gene-level visualization for longitudinally collected plasma samples for patient P6199.** Gene-level methylation is shown from the analysis of longitudinal cfDNA data as well as the matched tumor and immune methylomes. The top and bottom 25 genes with differing methylation between the primary tumor and immune cells were selected. Gray boxes indicate no reads were obtained for that sample.

### P4822 – Metastatic pancreatic neuroendocrine carcinoma

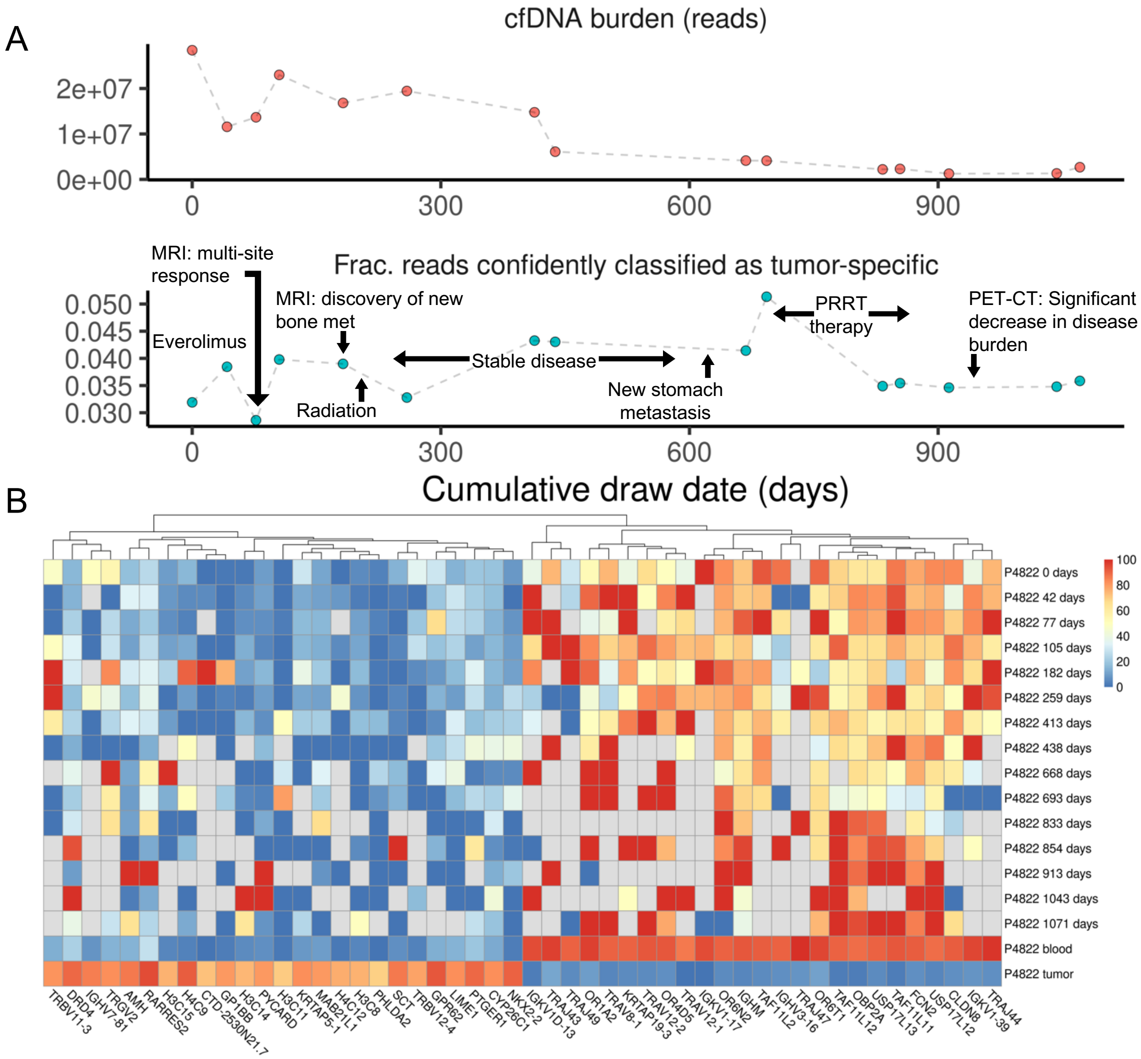


**Supplementary Figure 14. Longitudinal monitoring and gene-level visualization for patient P4822 with metastatic pancreatic neuroendocrine carcinoma. (A)** The overall cfDNA sequencing yield (top) is plotted against the number of reads with methylation profiles matching the primary tumor with a tumor score of >0.9 (bottom). Clinically relevant events are annotated. Clinically relevant events are annotated. Everolimus and peptide receptor radionuclide therapy (PRRT) was used for treatment of the metastatic neuroendocrine cancer. Positron-emission tomography (PET) was combined with CT imaging. **(B)** Gene-level methylation is shown from the analysis of longitudinal cfDNA data as well as the matched tumor and immune methylomes. The top and bottom 25 genes with differing methylation between the primary tumor and immune cells were selected. Gray boxes indicate no reads were obtained for that sample.

### P6527 – Intrahepatic cholangiocarcinoma

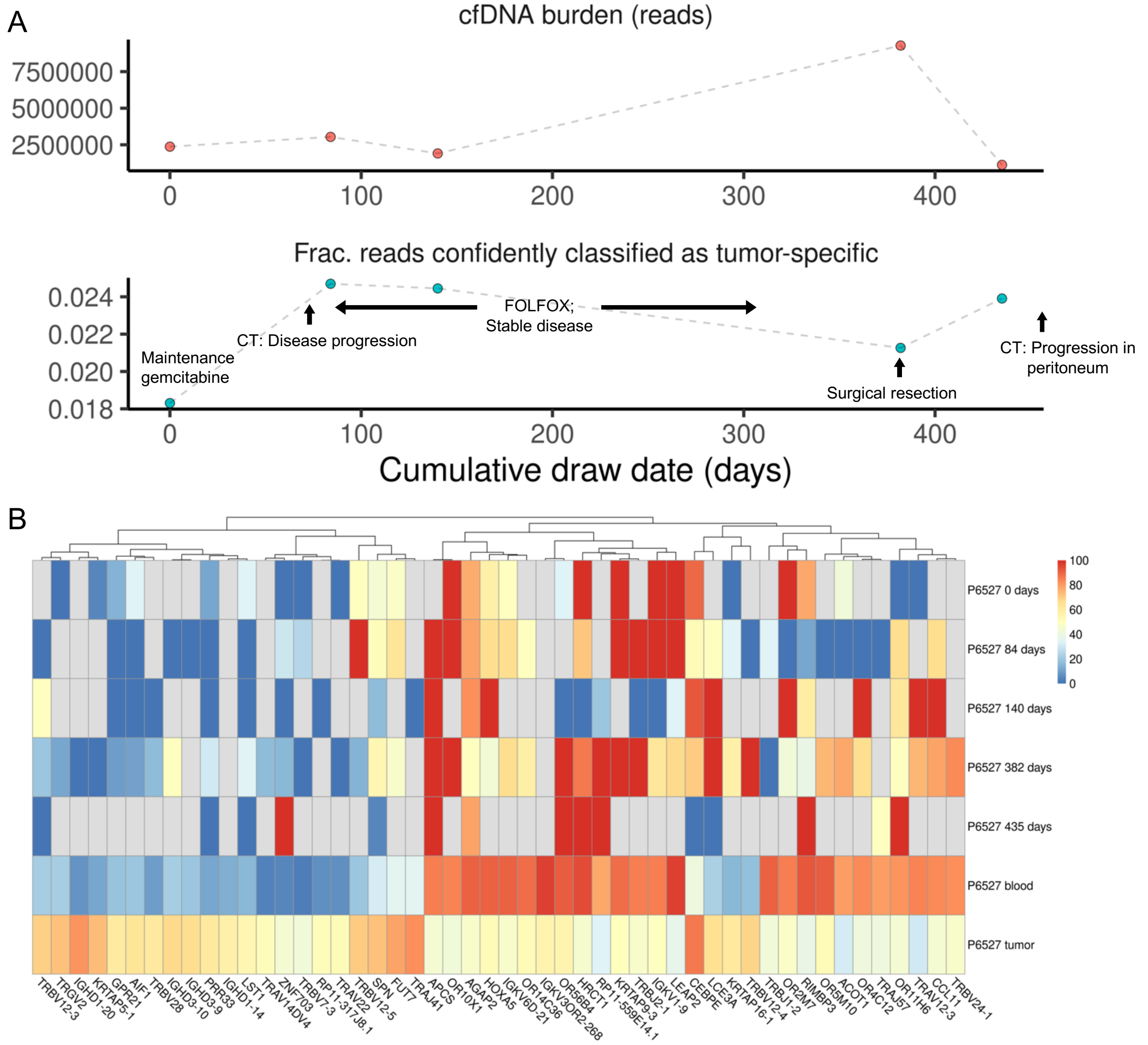


**Supplementary Figure 15. Longitudinal monitoring and gene-level visualization for patient P6527 with metastatic cholangiocarcinoma. (A)** The overall cfDNA sequencing yield (top) is plotted against the number of reads with methylation profiles matching the primary tumor with a tumor score of >0.9 (bottom). Clinically relevant events are annotated. Treatment included gemcitabine and a chemotherapy combination of 5-fluouracil and oxaliplatin (FOLFOX) was used for treatment of the cholangiocarcinoma. **(B)** Gene-level methylation is shown from the analysis of longitudinal cfDNA data as well as the matched tumor and immune methylomes. The top and bottom 25 genes with differing methylation between the primary tumor and immune cells were selected. Gray boxes indicate no reads were obtained for that sample.

**Supplementary Table 1. Sequencing Metrics**

| Patient | Sample | Experiment/Cohort | # reads | # aligned | % aligned | # CpG sites sequenced |
|---|---|---|---|---|---|---|
| P6604 | P6604_nucl | Protocol Comparison - This work - 5ng - replicate 1 | 7.34M | 6.47M | 88% | N/A |
| P6604 | P6604_nucl | Protocol Comparison - This work - 5ng - replicate 2 | 7.85M | 6.73M | 86% | N/A |
| P6604 | P6604_nucl | Protocol Comparison - This work - 5ng - replicate 3 | 9.49M | 4.70M | 50% | N/A |
| P6604 | P6604_nucl | Protocol Comparison - This work - 1ng - replicate 1 | 1.84M | 1.44M | 78% | N/A |
| P6604 | P6604_nucl | Protocol Comparison - This work - 1ng - replicate 2 | 1.61M | 1.31M | 82% | N/A |
| P6604 | P6604_nucl | Protocol Comparison - This work - 1ng - replicate 3 | 1.69M | 1.36M | 81% | N/A |
| P6604 | P6604_nucl | Protocol Comparison - This work - 0.5ng - replicate 1 | .89M | .71M | 79% | N/A |
| P6604 | P6604_nucl | Protocol Comparison - This work - 0.5ng - replicate 2 | 1.37M | .82M | 60% | N/A |
| P6604 | P6604_nucl | Protocol Comparison - This work - 0.5ng - replicate 3 | .81M | .63M | 77% | N/A |
| P6604 | P6604_nucl | Protocol Comparison - This work - 0.1ng - replicate 1 | .21M | .13M | 60% | N/A |
| P6604 | P6604_nucl | Protocol Comparison - This work - 0.1ng - replicate 2 | .37M | .16M | 42% | N/A |
| P6604 | P6604_nucl | Protocol Comparison - This work - 0.1ng - replicate 3 | .34M | .14M | 43% | N/A |
| P6604 | P6604_nucl | Protocol Comparison - ONT NBD196 protocol - 5ng - replicate 1 | 1.57M | 1.34M | 85% | N/A |
| P6604 | P6604_nucl | Protocol Comparison - ONT NBD196 protocol - 5ng - replicate 2 | 1.49M | 1.26M | 84% | N/A |
| P6604 | P6604_nucl | Protocol Comparison - ONT NBD196 protocol - 5ng - replicate 3 | 1.43M | 1.22M | 85% | N/A |
| P6604 | P6604_nucl | Protocol Comparison - ONT NBD196 protocol - 1ng - replicate 1 | .33M | .27M | 84% | N/A |
| P6604 | P6604_nucl | Protocol Comparison - ONT NBD196 protocol - 1ng - replicate 2 | .34M | .28M | 84% | N/A |
| P6604 | P6604_nucl | Protocol Comparison - ONT NBD196 protocol - 1ng - replicate 3 | .43M | .36M | 84% | N/A |
| P6604 | P6604_nucl | Protocol Comparison - ONT NBD196 protocol - 0.5ng - replicate 1 | .12M | .10M | 83% | N/A |
| P6604 | P6604_nucl | Protocol Comparison - ONT NBD196 protocol - 0.5ng - replicate 2 | .13M | .10M | 83% | N/A |
| P6604 | P6604_nucl | Protocol Comparison - ONT NBD196 protocol - 0.5ng - replicate 3 | .13M | .11M | 84% | N/A |
| P6604 | P6604_nucl | Protocol Comparison - ONT NBD196 protocol - 0.1ng - replicate 1 | .03M | .02M | 82% | N/A |
| P6604 | P6604_nucl | Protocol Comparison - ONT NBD196 protocol - 0.1ng - replicate 2 | .03M | .03M | 80% | N/A |
| P6604 | P6604_nucl | Protocol Comparison - ONT NBD196 protocol - 0.1ng - replicate 3 | .03M | .02M | 81% | N/A |
| GP2D | GP2D_nucl | In silico admixture | 97.06M | 90.64M | 93% | 56.41M |
| P6604 | P6604_nucl | In silico admixture | 115.41M | 101.09M | 88% | 42.62M |
| P6604/GP2D | P6604_nucl / GP2D_nucl | Experimental admixture - 500 pg - 100% GP2D | .41M | .11M | 26% | .15M |
| P6604/GP2D | P6604_nucl / GP2D_nucl | Experimental admixture - 500 pg - 50% GP2D | .64M | .18M | 29% | .16M |
| P6604/GP2D | P6604_nucl / GP2D_nucl | Experimental admixture - 500 pg - 10% GP2D | .51M | .23M | 44% | .17M |
| P6604/GP2D | P6604_nucl / GP2D_nucl | Experimental admixture - 500 pg - 5% GP2D | .66M | .27M | 40% | .19M |
| P6604/GP2D | P6604_nucl / GP2D_nucl | Experimental admixture - 250 pg - 100% GP2D | .33M | .05M | 16% | .08M |
| P6604/GP2D | P6604_nucl / GP2D_nucl | Experimental admixture - 250 pg - 50% GP2D | .28M | .08M | 29% | .08M |
| P6604/GP2D | P6604_nucl / GP2D_nucl | Experimental admixture - 250 pg - 10% GP2D | .44M | .12M | 26% | .08M |
| P6604/GP2D | P6604_nucl / GP2D_nucl | Experimental admixture - 250 pg - 5% GP2D | .42M | .12M | 29% | .09M |
| P6604/GP2D | P6604_nucl / GP2D_nucl | Experimental admixture - 125 pg - 100% GP2D | .46M | .03M | 7% | .04M |
| P6604/GP2D | P6604_nucl / GP2D_nucl | Experimental admixture - 125 pg - 50% GP2D | .34M | .04M | 13% | .04M |
| P6604/GP2D | P6604_nucl / GP2D_nucl | Experimental admixture - 125 pg - 10% GP2D | .48M | .06M | 12% | .04M |
| P6604/GP2D | P6604_nucl / GP2D_nucl | Experimental admixture - 125 pg - 5% GP2D | .48M | .06M | 13% | .05M |
| P6604/GP2D | P6604_nucl / GP2D_nucl | Experimental admixture - 63 pg - 100% GP2D | .27M | .02M | 8% | .03M |
| P6604/GP2D | P6604_nucl / GP2D_nucl | Experimental admixture - 63 pg - 50% GP2D | .41M | .04M | 10% | .04M |
| P6604/GP2D | P6604_nucl / GP2D_nucl | Experimental admixture - 63 pg - 10% GP2D | .45M | .05M | 11% | .04M |
| P6604/GP2D | P6604_nucl / GP2D_nucl | Experimental admixture - 63 pg - 5% GP2D | .15M | .05M | 32% | .04M |
| P6600 | P6600_plas | 20 patient cohort - Healthy Control | 3.01M | 1.37M | 46% | 3.02M |
| P6601 | P6601_plas | 20 patient cohort - Healthy Control | 4.56M | 1.69M | 37% | 3.71M |
| P6602 | P6602_plas | 20 patient cohort - Healthy Control | 6.97M | 2.10M | 30% | 3.12M |
| P6603 | P6603_plas | 20 patient cohort - Healthy Control | 2.64M | .66M | 25% | 1.77M |
| P2574 | P2574_3331A | 20 patient cohort - Cancer | 16.42M | 14.54M | 89% | 11.66M |
| P2592 | P2592_3329A | 20 patient cohort - Cancer | 2.19M | 1.95M | 89% | 2.07M |
| P3621 | P3621_3317A | 20 patient cohort - Cancer | 1.89M | 1.68M | 89% | 2.26M |
| P4776 | P4776_3319A | 20 patient cohort - Cancer | 3.71M | 3.37M | 91% | 3.41M |
| P5070 | P5070_3339A | 20 patient cohort - Cancer | 197.66M | 179.80M | 91% | 52.48M |
| P741 | P741_3303A | 20 patient cohort - Cancer | 1.67M | 1.26M | 76% | 2.39M |
| P9075 | P9075_3301A | 20 patient cohort - Cancer | 1.60M | 1.40M | 88% | 1.69M |
| P9076 | P9076_3305A | 20 patient cohort - Cancer | 2.59M | 2.37M | 91% | 2.94M |
| P9077 | P9077_3307A | 20 patient cohort - Cancer | 72.23M | 64.74M | 90% | 33.69M |
| P9078 | P9078_3309A | 20 patient cohort - Cancer | 59.52M | 54.30M | 91% | 35.20M |
| P9079 | P9079_3311A | 20 patient cohort - Cancer | 2.01M | 1.71M | 85% | 2.11M |
| P9080 | P9080_3313A | 20 patient cohort - Cancer | 10.37M | 9.45M | 91% | 8.83M |
| P9081 | P9081_3315A | 20 patient cohort - Cancer | 2.11M | 1.63M | 77% | 2.14M |
| P9082 | P9082_3321A | 20 patient cohort - Cancer | 13.85M | 12.40M | 90% | 10.34M |
| P9083 | P9083_3323A | 20 patient cohort - Cancer | 52.56M | 47.41M | 90% | 30.97M |
| P9084 | P9084_3325A | 20 patient cohort - Cancer | 2.82M | 2.47M | 88% | 3.35M |
| P9085 | P9085_3327A | 20 patient cohort - Cancer | 8.91M | 7.88M | 88% | 6.78M |
| P9086 | P9086_3333A | 20 patient cohort - Cancer | .81M | .67M | 83% | 2.64M |
| P9087 | P9087_3335A | 20 patient cohort - Cancer | 1.35M | 1.20M | 89% | 2.11M |
| P9088 | P9088_3337A | 20 patient cohort - Cancer | 54.10M | 49.21M | 91% | 29.96M |
| P6199 | P6199_19520 | Longitudinal | 25.65M | 22.92M | 89% | 17.18M |
| P6199 | P6199_19526 | Longitudinal | 11.40M | 10.20M | 90% | 7.72M |
| P6199 | P6199_19572 | Longitudinal | 17.67M | 15.86M | 90% | 13.55M |
| P6199 | P6199_19585 | Longitudinal | 9.10M | 8.08M | 89% | 6.25M |
| P6199 | P6199_19620 | Longitudinal | 9.89M | 8.93M | 90% | 8.85M |
| P6199 | P6199_21306 | Longitudinal | 27.27M | 24.86M | 91% | 18.69M |
| P6199 | P6199_21347 | Longitudinal | 6.03M | 5.46M | 91% | 6.46M |
| P6199 | P6199_21381 | Longitudinal | 13.33M | 12.02M | 90% | 12.43M |
| P6199 | P6199_21433 | Longitudinal | 11.91M | 10.67M | 90% | 9.70M |
| P6199 | P6199_21465 | Longitudinal | 9.46M | 8.47M | 90% | 8.28M |
| P6199 | P6199_21488 | Longitudinal | 10.23M | 9.32M | 91% | 9.63M |
| P6199 | P6199_21536 | Longitudinal | 4.87M | 4.18M | 86% | 4.51M |
| P6199 | P6199_21549 | Longitudinal | 6.02M | 5.41M | 90% | 6.44M |
| P6199 | P6199_21641 | Longitudinal | 3.13M | 2.67M | 85% | 2.88M |
| P6199 | P6199_21671 | Longitudinal | 7.70M | 6.87M | 89% | 7.08M |
| P6199 | P6199_immune | Longitudinal | 14.94M | 14.65M | 98% | 56.86M |
| P6199 | P6199_primary | Longitudinal | 29.43M | 29.04M | 99% | 57.99M |
| P4822 | P4822_18277 | Longitudinal | 57.12M | 51.23M | 90% | 30.81M |
| P4822 | P4822_18284 | Longitudinal | 23.61M | 20.94M | 89% | 17.07M |
| P4822 | P4822_18309 | Longitudinal | 26.81M | 24.14M | 90% | 19.17M |
| P4822 | P4822_18317 | Longitudinal | 46.14M | 41.39M | 90% | 27.94M |
| P4822 | P4822_18404 | Longitudinal | 33.10M | 29.69M | 90% | 22.19M |
| P4822 | P4822_18452 | Longitudinal | 37.91M | 33.91M | 89% | 24.78M |
| P4822 | P4822_19469 | Longitudinal | 30.17M | 26.79M | 89% | 18.89M |
| P4822 | P4822_19502 | Longitudinal | 12.90M | 11.51M | 89% | 8.03M |
| P4822 | P4822_21383 | Longitudinal | 8.46M | 7.60M | 90% | 6.03M |
| P4822 | P4822_21431 | Longitudinal | 8.28M | 7.24M | 87% | 7.13M |
| P4822 | P4822_21518 | Longitudinal | 4.35M | 3.87M | 89% | 3.97M |
| P4822 | P4822_21540 | Longitudinal | 4.64M | 4.02M | 87% | 4.21M |
| P4822 | P4822_21581 | Longitudinal | 2.52M | 2.26M | 90% | 2.05M |
| P4822 | P4822_21682 | Longitudinal | 2.55M | 2.24M | 88% | 2.50M |
| P4822 | P4822_22186 | Longitudinal | 4.93M | 4.44M | 90% | 5.38M |
| P4822 | P4822_immune | Longitudinal | 13.08M | 12.90M | 99% | 57.90M |
| P4822 | P4822_primary | Longitudinal | 27.39M | 26.97M | 98% | 57.70M |
| P6527 | P6527_19633 | Longitudinal | 4.71M | 3.82M | 81% | 5.29M |
| P6527 | P6527_21349 | Longitudinal | 6.43M | 5.05M | 79% | 6.83M |
| P6527 | P6527_21416 | Longitudinal | 4.45M | 3.31M | 74% | 3.75M |
| P6527 | P6527_21602 | Longitudinal | 16.68M | 14.96M | 90% | 18.36M |
| P6527 | P6527_21639 | Longitudinal | 2.78M | 1.90M | 68% | 2.38M |
| P6527 | P6527_immune | Longitudinal | 17.30M | 16.99M | 98% | 58.23M |
| P6527 | P6527_primary | Longitudinal | 31.53M | 31.07M | 99% | 58.27M |

**Supplementary Table 2. Patient Information**

| Patient | Reported Diagnosis | TNM staging reported at surgery | Number of time points | Primary tumor available/sequenced |
|---|---|---|---|---|
| P4822 | Metastatic Pancreatic Neuroendocrine Carcinoma | N/A | 14 | Y |
| P6199 | Invasive Adenocarcinoma, Poorly Differentiated, Extending Into Pericolonic Soft Tissue | ypT3 pN2b | 15 | Y |
| P6527 | Intrahepatic Cholangiocarcinoma, Moderately Differentiated | ypT2 pN0 | 5 | Y |
| P9075 | Invasive Adenocarcinoma | pT3 pN2a | 1 | N |
| P741 | Moderately Differentiated Colorectal Adenocarcinoma | pT4 pN2 pM1 | 1 | N |
| P9076 | Recurrent Adenocarcinoma, Colorectal Primary | pT3 pN1a | 1 | N |
| P9077 | Invasive Adenocarcinoma | pT3 pN1b pM1a | 1 | N |
| P9078 | Invasive Adenocarcinoma | ypT2 ypN0 ypM0 | 1 | N |
| P9079 | Adenocarcinoma, Cribriform Comedo Type | pT2 pN0 | 1 | N |
| P9080 | Invasive Adenocarcinoma/Metastatic Adenocarcinoma | ypT2 ypN2a ypM1 | 1 | N |
| P9081 | Invasive Adenocarcinoma | ypT4b ypN0 | 1 | N |
| P3621 | Invasive Colorectal Adenocarcinoma | pT3 pN0 | 1 | N |
| P4776 | Invasive Adenocarcinoma | pT4b pN1b | 1 | N |
| P9082 | Metastatic Adenocarcinoma, Colorectal Primary | pT4b pN1b | 1 | N |
| P9083 | Metastatic Adenocarcinoma | N/A | 1 | N |
| P9084 | Metastatic Adenocarcinoma, Colorectal Primary | N/A | 1 | N |
| P9085 | Metastatic Adenocarcinoma, Colorectal Primary | ypT4b pN0 pM1a | 1 | N |
| P2592 | Metastatic Adenocarcinoma In Three Of Four Lymph Nodes | N/A | 1 | N |
| P2574 | Metastatic Adenocarcinoma, Colorectal Primary | T3N0 | 1 | N |
| P9086 | Metastatic Adenocarcinoma,Colorectal Primary | ypT1 pN1c pM1a | 1 | N |
| P9087 | Metastatic Adenocarcinoma, Colonic Primary | pT3 pN1a | 1 | N |
| P9088 | Metastatic Adenocarcinoma, Colorectal Primary | N/A | 1 | N |
| P5070 | Metastatic Adenocarcinoma | N/A | 1 | N |

**Supplementary Table 3. Genes with significant methylation differences between healthy and patient-derived cfDNA in 20 patient cohort**

| Gene | q-value (fdr adjusted) | Mean difference between groups (healthy - cancer) |
|---|---|---|
| SPIB | 3.70E-06 | -41.39933674 |
| CDCA7 | 6.52E-06 | -53.22244908 |
| TMEM164 | 2.36E-05 | 18.216444 |
| COL10A1 | 0.00015059 | 17.13642991 |
| PLSCR4 | 0.000182298 | -34.87794316 |
| SLC25A1 | 0.000182298 | 60.89159323 |
| ELAC2 | 0.000447691 | 25.31557705 |
| ZNF572 | 0.000503628 | -59.14379304 |
| ENPP4 | 0.000600976 | 45.02995548 |
| GPER1 | 0.000616948 | 26.90489456 |
| PLAGL2 | 0.000616948 | 37.94148816 |
| RPUSD4 | 0.000616948 | 37.03039294 |
| NUF2 | 0.000836028 | -20.19624653 |
| SMIM10L2A | 0.00083887 | 54.84387916 |
| GJC1 | 0.001031567 | 33.99002109 |
| ILRUN | 0.001163916 | 10.93166634 |
| ZNF414 | 0.001198392 | -60.93005356 |
| ZNF772 | 0.001198392 | 35.83525929 |
| ELL2 | 0.00154547 | -30.51454594 |
| KLHL11 | 0.00154547 | 33.69456263 |
| LGR4 | 0.00160918 | -20.65861339 |
| LHCGR | 0.001730388 | 20.15967146 |
| ADGRG4 | 0.002000412 | 29.8121374 |
| CTD-2545M3.6 | 0.002000412 | -41.10365785 |
| ELAPOR2 | 0.002000412 | 17.26097167 |
| MGMT | 0.002000412 | 9.027538374 |
| NME1 | 0.002177991 | -51.33222625 |
| NRTN | 0.002240827 | 20.54819725 |
| OSGEPL1 | 0.002240827 | 33.21504636 |
| AMDHD1 | 0.002413003 | 36.1677824 |
| MCHR2 | 0.002413003 | 25.94704782 |
| ROGDI | 0.002413003 | 43.62026154 |
| ZNF774 | 0.002664044 | 33.96611201 |
| RER1 | 0.003040081 | 26.67522645 |
| OPN1MW2 | 0.00313277 | 29.30895359 |
| TUG1 | 0.00313277 | 44.06717689 |
| CAPN11 | 0.003149901 | 24.94768262 |
| MRPL52 | 0.003149901 | -40.1475143 |
| KIAA1143 | 0.003187013 | 24.34567563 |
| INTS6L | 0.00397628 | 27.17036478 |
| ARMCX5-GPRASP2 | 0.004016938 | 29.95589663 |
| PSMD2 | 0.004016938 | 35.58255254 |
| PUS3 | 0.004285724 | 43.49319279 |
| LAMP2 | 0.004563378 | 22.05689646 |
| TBC1D22A | 0.004563378 | 5.681992433 |
| TRIM51 | 0.004563378 | 43.78787467 |
| UCN | 0.004563378 | 73.24983018 |
| CD79A | 0.005618985 | -48.92757848 |
| DNHD1 | 0.005909211 | 11.01400976 |
| ADAMTS14 | 0.006008943 | 12.90419821 |
| RNA5S5 | 0.006008943 | 51.83139542 |
| ULBP3 | 0.006008943 | 33.96751199 |
| EPPIN | 0.006158961 | 44.53709195 |
| SSU72 | 0.006158961 | 20.63218936 |
| ZBTB22 | 0.006220455 | -38.11904119 |
| FAIM | 0.006354143 | 34.27565271 |
| TCTA | 0.006503272 | 52.32468329 |
| OR4D2 | 0.006583676 | 39.60481249 |
| RETSAT | 0.006583676 | 32.24024626 |
| RNFT1 | 0.006583676 | -46.78586976 |
| HSD3B7 | 0.006606003 | 36.44872641 |
| NXF2 | 0.006695395 | 17.65769516 |
| DICER1 | 0.007310389 | 14.5453317 |
| CNKSR2 | 0.007376806 | 22.62391933 |
| MAGEA9B | 0.007376806 | 32.9689437 |
| GSTT2B | 0.007446381 | 42.19318471 |
| TCL1B | 0.007446381 | 30.32851384 |
| KAT2A | 0.007576995 | 35.28736586 |
| NFE2L3 | 0.007683316 | 22.96929258 |
| TMEM150A | 0.008110172 | 50.78549013 |
| ZIC3 | 0.008110172 | -14.66498911 |
| LAMTOR4 | 0.008516659 | 34.7124756 |
| CSDE1 | 0.008686958 | -23.89282634 |
| MRPS18A | 0.008952295 | 27.97364516 |
| CCDC71 | 0.009340928 | 43.7421799 |
| CCL16 | 0.009555689 | 33.05517035 |
| LAMTOR2 | 0.009555689 | 48.04111602 |
| OR13A1 | 0.009555689 | 34.48973445 |
| GPATCH4 | 0.009717914 | -44.38019697 |
| EPS8L3 | 0.009912777 | 15.53967346 |

**Supplementary Table 4. Methods Comparison**

| Method | Sequencing Platform | Resolution | PCR-free | Input requirement | Comments |
|---|---|---|---|---|---|
| This work | Oxford Nanopore | Base-pair | Yes | pg to ng | Utlizes LSK110 latest chemistry on R9.4.1 flow cells |
| Conventional Nanopore | Oxford Nanopore | Base-pair | Yes | >40ng | Barcoding adapters are stuck with an previous generation sequencing adapter |
| Bisulfite | llumina | Base-pair | No | tens to hundreds of ng | |
| Enzymatic | llumina | Base-pair | No | tens to hundreds of ng | |
| cfMeDIP-Seq | Illumina | Binned | Yes | ng to hundreds of ng | requires carrier |
